# Structural modeling reveals the allosteric switch controlling the chitin utilization program of *Vibrio cholerae*

**DOI:** 10.1101/2025.04.22.650076

**Authors:** Holly S. Anderson, Ankur B. Dalia

## Abstract

Signal transduction by histidine kinases (HKs) is nearly ubiquitous in bacterial species. HKs can either sense ligands directly or indirectly via a cognate solute binding protein (SBP). The molecular basis for SBP-dependent signal reception, however, remains poorly understood in most cases. CBP and ChiS are the SBP-HK pair that activate the chitin utilization program of *Vibrio cholerae*. Here, we elucidate the molecular basis for allosteric regulation of CBP-ChiS by generating structural models of this complex in the unliganded and liganded states, which we support with extensive genetic, biochemical, and cell biological analysis. Our results reveal that ligand-binding induces a large conformational interface switch that is distinct from previously described SBP-HKs. Structural modeling suggests that similar interface switches may also regulate other uncharacterized SBP-HKs. Together, these results extend our understanding of signal transduction in bacterial species and highlight a new approach for uncovering the molecular basis of allostery in protein complexes.

**SIGNIFICANCE STATEMENT:** All living things use protein receptors to sense and respond to environmental changes via a process termed signal transduction. However, how these proteins sense environmental stimuli remains poorly understood in many cases. In this study, we study allosteric activation of the chitin sensor, ChiS, by chitin-binding protein, CBP, in *Vibrio cholerae* as a model system. Using a combination of structural modeling, genetics, and biochemistry we uncovered the molecular basis underlying CBP-ChiS allosteric regulation, which we find is distinct from previously described systems. This work expands our understanding of bacterial signal transduction and highlights an approach for uncovering new modes of allosteric regulation.

## INTRODUCTION

The ability to sense and respond appropriately to environmental stimuli is critical in all domains of life. One sensory system that is nearly ubiquitous in bacterial species is signal transduction by histidine kinases (HKs)^1,2^. HKs can sense periplasmic signals either directly^3^ or indirectly via a solute binding protein (SBP)^4^. While genetic studies have linked SBPs to activation of cognate HKs^4–8^, the molecular mechanism for activation remains poorly understood in most cases. This is due, in part, to the challenges associated with structural characterization of transmembrane proteins like HKs. As a result, the current understanding of SBP-HK activation is informed by a relatively limited number of structural studies^9–12^. This work has revealed that the molecular basis of SBP-HK interactions differs greatly between systems, indicating the lack of a broadly conserved allosteric activation mechanism.

While activation of HKs by liganded SBPs is well-studied, the influence of unliganded SBPs on HK regulation has been overlooked in most cases. Paradigmatically, unliganded SBPs interact passively with their cognate HKs and then allosterically induce activation through a conformational change upon ligand-binding^9–12^. The <u>chi</u>tin-<u>s</u>ensing HK ChiS offers a unique and unexplored example of SBP-HK regulation. ChiS is a transmembrane transcriptional regulator that directly binds DNA^6^ to regulate the expression of genes required for chitin catabolism and natural transformation in *V. cholerae* in response to chitin^13–15^. ChiS has both kinase and phosphatase activity, and the phosphorylation state of this one-component HK regulates its DNA-binding activity^6^. Prior genetic and cell biological analysis indicates that ChiS is subject to dual allosteric regulation by the periplasmic SBP, <u>c</u>hitin-<u>b</u>inding <u>p</u>rotein (CBP)^6^. In the absence of chitin, unliganded CBP represses ChiS DNA-binding activity in a manner that is dependent on ChiS kinase activity^6^. Conversely, liganded (chitin-bound) CBP interacts with ChiS to inhibit its kinase activity, resulting in dephosphorylation of ChiS and induction of its DNA-binding activity^6^. Here, we sought to define the molecular basis underlying this dual allosteric regulation to expand our understanding of bacterial signal transduction.

## RESULTS

### AlphaFold predicts the CBP-ChiS complex in two distinct conformations

To help define the molecular basis for CBP-dependent allosteric regulation of ChiS activity, we sought to generate structural models of CBP-ChiS using AlphaFold-Multimer v3 (AF-Mv3)^16–18^ and AlphaFold3 (AF3)^19^. Because HKs canonically form homodimers^1^, we modeled two copies of ChiS, and we empirically found that structural modeling with one CBP yielded models of the highest confidence and with the fewest steric clashes for both algorithms (**Fig. S1**). AF-Mv3 and AF3 modeled ChiS as a homodimer, as expected, with CBP cradled at its apex (**Fig. S2**). Interestingly, the CBP-ChiS^AF-M^ and CBP-ChiS^AF3^ models predicted distinct orientations for CBP (**Fig. S2, Movie S1**) despite having similar confidence scores (**Fig. S1A-B**). Since CBP differentially regulates ChiS activity depending on its liganded state^6^, we hypothesized that these two CBP-ChiS models represented the unliganded (ChiS repressed) and liganded (ChiS active) conformations. To test this hypothesis, we aligned CBP from the top-ranked CBP-ChiS^AF-M^ and CBP-ChiS^AF3^ models to crystal structures of unliganded (PDB: 4gf8) and liganded (PBD: 4gfr) CBP. This analysis revealed that CBP^AF-M^ closely aligned to liganded CBP, while CBP^AF3^ aligned to unliganded CBP (**Fig. S3**). Thus, we hypothesized that the CBP-ChiS^AF-M^ model represents the liganded (chitin-bound) conformation where ChiS is active, while the CBP-ChiS^AF3^ model represents the unliganded conformation where ChiS is repressed (**Fig. 1**).

**Fig. 1.**
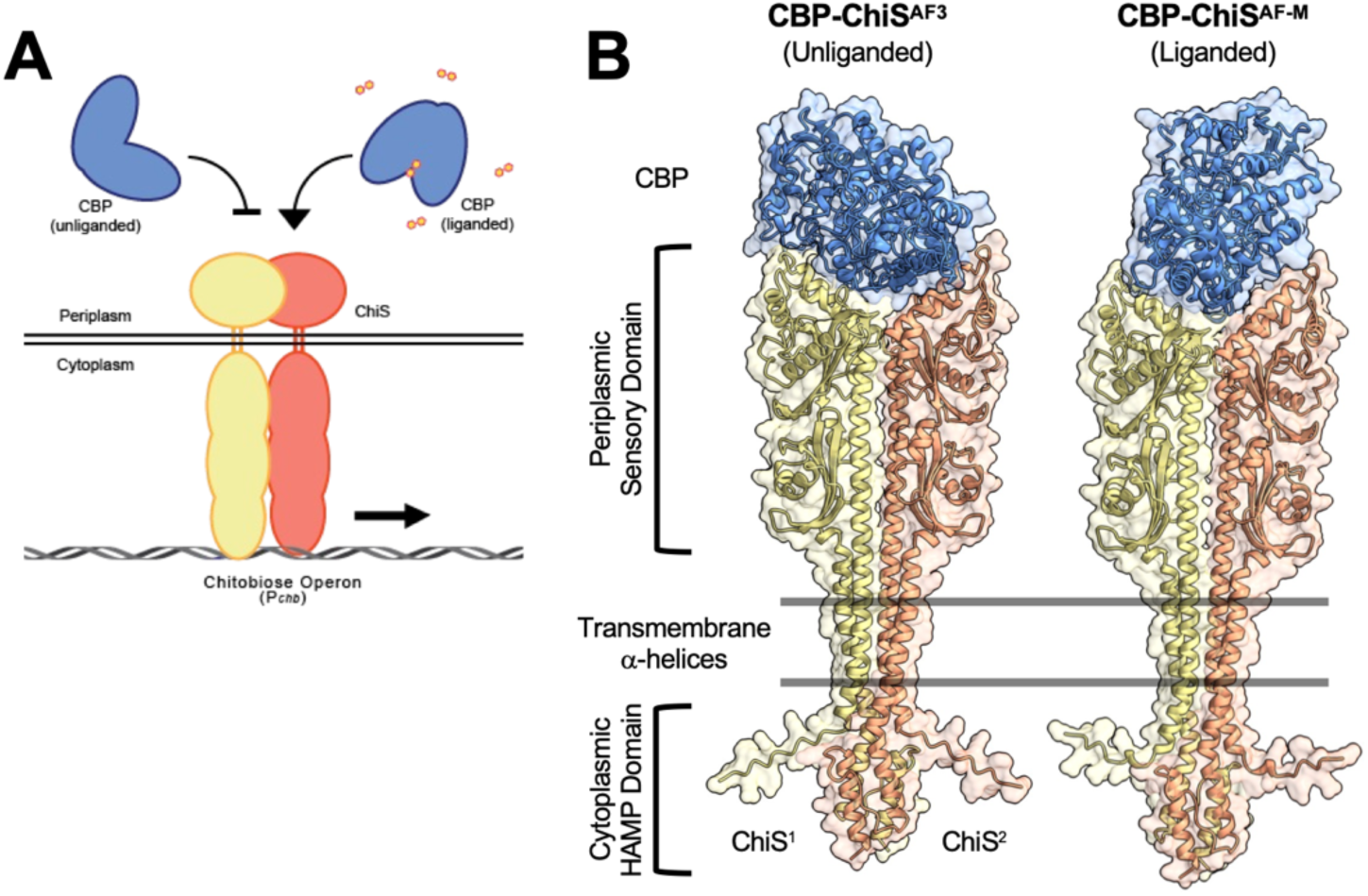
Structural modeling of CBP-ChiS reveals distinct conformations. (**A**) Schematic representation of CBP dual allosteric regulation of ChiS activity. Unliganded CBP represses ChiS activity, while liganded CBP stimulates ChiS activity. (**B**) Top ranked models of CBP-ChiS^AF3^ and CBP-ChiS^AF-M^. CBP is colored blue, while the two copies of the ChiS periplasmic domain (residues 1-427) are colored yellow & orange.

### Comparison of the CBP-ChiS^AF3^ and CBP-ChiS^AF-M^ models reveals interfacial salt bridges unique to each model

To formally test this hypothesis, we sought to identify interactions unique to each model for targeted mutagenesis. We focused on identifying residue pairs that putatively form interfacial salt bridges because they allow for a high degree of specificity in structure-function analysis. Specifically, mutating these residues to the opposite charge should break interactions through electrostatic repulsion. However, if these residues truly interact, flipping the charge on both sides of the interface should restore the interaction. Thus, we analyzed the top-ranked models from five independent CBP-ChiS^AF-M^ and CBP-ChiS^AF3^ sessions to identify interfacial salt bridges at the CBP-ChiS interface. Interactions predicted in ≥3 models that were separated by <4 Å in one conformation but >10 Å in the other conformation (**Fig. S4A, Dataset S1**) were selected for further characterization. This analysis revealed one intermolecular salt bridge specific to the CBP-ChiS^AF3^ model (unliganded / repressed conformation) and five specific to the CBP-ChiS^AF-M^ model (liganded / active conformation) (**Fig. S4B-C**).

### Interactions specific to the CBP-ChiS^AF3^ (unliganded) model are critical for CBP to allosterically repress ChiS in the absence of chitin

In the absence of chitin, unliganded CBP interacts with ChiS to allosterically repress its activity^6,15^. We hypothesized that interactions specific to the CBP-ChiS^AF3^ model (**Fig. 2A, S4**) are necessary for CBP-dependent repression of ChiS in the absence of chitin. Thus, flipping the charge of residues unique to the CBP-ChiS^AF3^ interface should disrupt CBP’s ability to repress ChiS, resulting in increased ChiS activity in rich media (*i.e.*, absence of chitin). Since CBP is directly regulated by ChiS^6^, a functional CBP-mCherry fusion under the control of its native promoter (P*_chb_*-*cbp*-*mCherry*) is used as a reporter for ChiS activity throughout this manuscript. The mCherry fluorescence is normalized to a constitutively expressed mTFP1 construct (P*_const2_-mTFP1*) to control for intrinsic noise in gene expression. Consistent with unliganded CBP repressing ChiS in the absence of chitin, ChiS activity is low when CBP is intact and inactivation of CBP (CBP^Δ^) results in elevated ChiS activity (**Fig. 2B**).

**Fig. 2.**
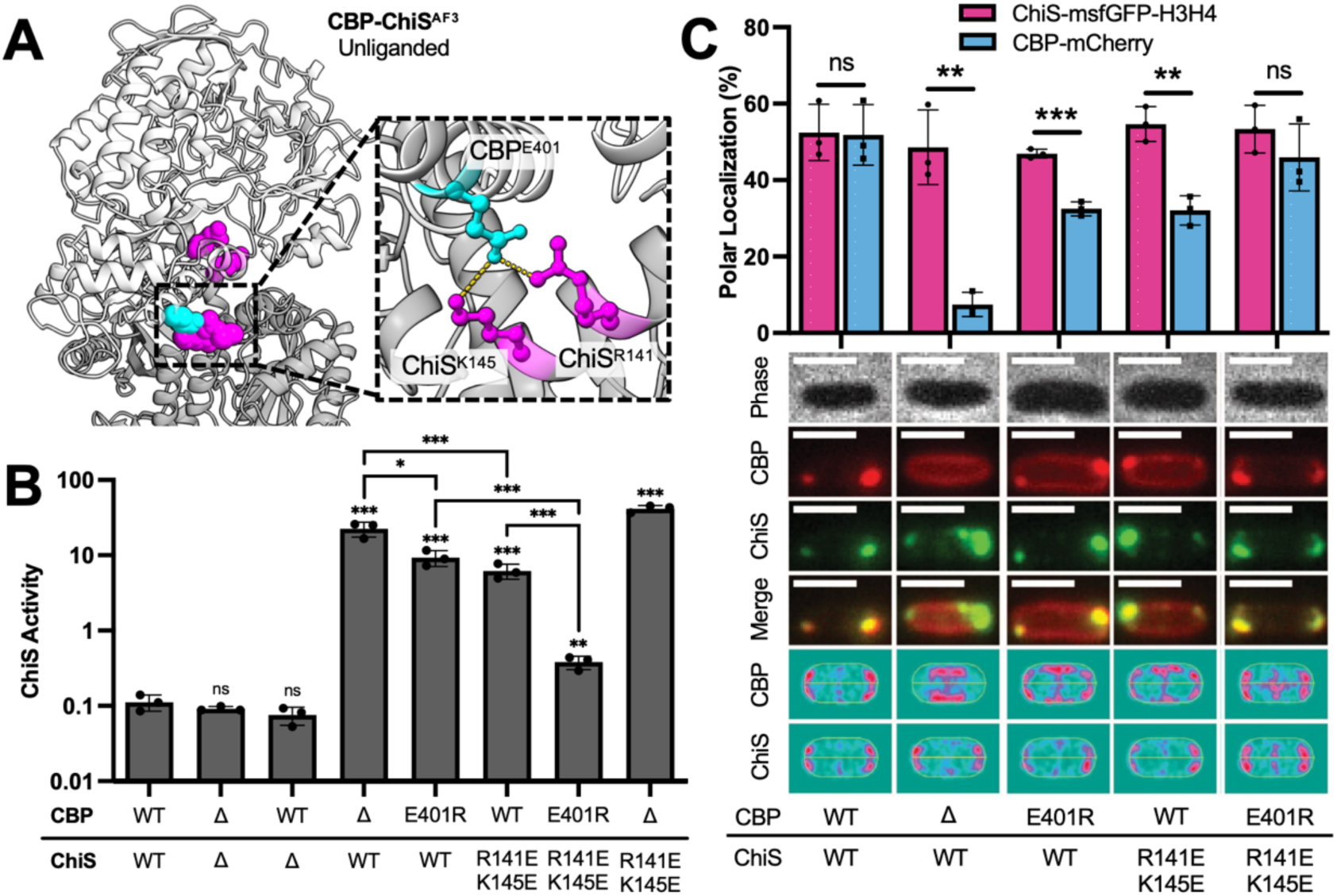
Intermolecular salt-bridges specific to CBP-ChiS^AF3^ (unliganded) is necessary for allosteric repression of ChiS in the absence of chitin. (**A**) Positions of CBP^E401^ (cyan) and ChiS^R141,K145^ (magenta) are highlighted in the CBP-ChiS^AF3^ (unliganded) model to show the intermolecular salt bridge (yellow dashed line) formed. CBP is shown in light grey, while ChiS is shown in dark grey. (**B**) Transcriptional reporter assays to measure ChiS activity in strains harboring the indicated mutations. All strains harbored P_chb_-cbp-mCherry and P_const2_-mTFP1 reporter constructs. The normalized mCherry fluorescence (see **Methods** for details) is reported as ChiS activity. Cells were grown in rich medium (i.e., in the absence of chitin) and fluorescence was determined by microscopy. Data are from three independent biological replicates (n = 300 cells analyzed per replicate) and shown as the mean ± SD. Statistical comparisons were made by one-way ANOVA with Tukey’s multiple comparison test on the log-transformed data. Statistical identifiers directly above bars represent comparisons to the parent. (**C**) CBP-ChiS interactions were assessed by POLAR assays. Cells contained *cbp-mCherry* and *chiS-msfGFP-H3H4* constructs with the indicated mutations as well as an inducible PopZ construct (P*_BAD_-popZ*). CBP^Δ^ = the CBP signal sequence translationally fused to mCherry (P*_chb_-cbp^1-40^-mCherry)*. Fluorescence microscopy was performed on cells with PopZ induction. Representative images are shown. Scale bars, 2 µm. Heat maps represent the localization of mCherry and GFP fluorescence in 300 cells. The proportion of CBP-mCherry (cyan bars) and ChiS-msfGFP-H3H4 (purple bars) fluorescence within the polar region (≤0.2 µm from the cell pole) was quantified. For each biological replicate (*n* = 3), fluorescence localization was determined by analyzing 300 individual cells. Statistical comparisons were made by Student’s t-test on the log-transformed data. ns = not significant, * = *p* < 0.05, ** = *p* < 0.01, *** = *p* < 0.001.

Mutating either side of the intermolecular salt bridge specific to CBP-ChiS^AF3^ (unliganded) (*cbp^E401R^* or *chiS^R141E,K145E^*) results in a significant increase in ChiS activity (**Fig. 2A-B**), which is consistent with these residues playing an important role in CBP-dependent repression of ChiS. Importantly, these mutations do not affect CBP or ChiS expression (**Fig. S5**) and the *chiS^R141E,K145E^* mutation does not diminish ChiS activity (**Fig. 2B**). To test whether these residues interact, we flipped the charge on both sides of the interface to restore the salt bridge. In *cbp^E401R^ chiS^R141E,K145E^*, CBP-dependent repression of ChiS is almost completely restored (**Fig. 2B**). These data strongly support the hypothesis that CBP^E401^-ChiS^R141,K145^ form a salt bridge and that CBP-ChiS^AF3^ represents the unliganded (repressed) complex conformation.

### An interfacial salt bridge specific to CBP-ChiS^AF3^ (unliganded) contributes to CBP-ChiS interactions

Unliganded CBP directly interacts with the periplasmic domain of ChiS to repress its activity^6^. Thus, we hypothesized that mutating the CBP^E401^-ChiS^R141,K145^ interface might prevent CBP-dependent repression by disrupting CBP-ChiS interactions. To test this, we used PopZ-linked apical recruitment (POLAR) assays^20^ to assess CBP-ChiS interactions by fluorescence microscopy^6^. In these assays, the “bait” and “prey” proteins are tagged with a fluorescent protein and the bait also contains an H3H4 PopZ-interaction domain. Upon induction of PopZ, the bait (ChiS-msfGFP-H3H4) relocalizes to the cell pole and the coincident relocalization of the prey (CBP-mCherry) demonstrates a direct interaction^20^ (**Fig. S6A**). Consistent with CBP-ChiS interactions, we see that CBP-mCherry and ChiS-msfGFP-H3H4 colocalize (**Fig. 2C, S6B**) and are both relocalized to the cell pole when PopZ is induced (**Fig. 2C**). A truncated allele of CBP (CBP^Δ^ = CBP signal sequence-mCherry fusion) does not interact with ChiS and remains peripherally localized, consistent with its periplasmic localization (**Fig. 2C, S6B**). When we flip the charge on either side of the putative unliganded interface (*cbp^E401R^*or *chiS^R141E,K145E^*), CBP-ChiS interactions are partially disrupted (**Fig. 2C, S6B**). Importantly, CBP-ChiS interactions are restored when the charge is flipped on both sides of the interface (*cbp^E401R^ chiS^R141E,K145E^*), which is consistent with restoration of the interfacial salt bridge (**Fig. 2C, S6B)**. Together, these results suggest that the salt bridge formed by CBP^E401^-ChiS^R141,K145^ is necessary for CBP-dependent repression in the absence of chitin because it contributes to CBP-ChiS interactions.

### Interactions specific to the ChiS^AF-M^ (liganded) model are necessary for CBP to allosterically activate ChiS in the presence of chitin

If CBP-ChiS^AF-M^ represents the liganded (active) conformation, we hypothesize that interactions unique to this model (**Fig. S4**) should be dispensable for CBP-dependent repression of ChiS in the absence of chitin. Consistent with this, we found that flipping the charge of residues unique to the CBP-ChiS^AF-M^ interface (*cbp^D231R^*, *cbp^D251R^*, *cbp^R254E^*, *cbp^E529R,E530R^*, *chiS^E81R^*, *chiS^R89E^*, *chiS^R99D^*, *chiS^R146D^*) did not impact CBP-dependent repression of ChiS in the absence of chitin with one exception (*chiS*^R139E^), which partially disrupted repression (**Fig. S7**). Thus, mutations to the CBP-ChiS^AF-M^ interface do not disrupt CBP-ChiS interactions in the unliganded state.

In the presence of chitin, liganded CBP allosterically activates ChiS activity^6^. If CBP-ChiS^AF-M^ represents the liganded (active) conformation, then mutating interactions specific to this interface should prevent chitin-dependent activation of ChiS. One potentially confounding issue in testing this hypothesis is that any disruption to CBP-ChiS interactions inherently increases ChiS activity, even in the absence of chitin (**Fig. 2B**). Contrary to most HKs, ChiS is active when it is unphosphorylated, and prior work demonstrates that liganded CBP allosterically activates ChiS by inhibiting its kinase activity^6^. Thus, a ChiS phosphatase-deficient mutant (*chiS^T473A^*) can only be activated in the presence of both CBP and chitin^6^, a finding that we recapitulate here (**Fig. S8**). Additionally, mutations to the CBP ligand-binding pocket (*cbp^W389A,W539A^*), that inhibit chitin-binding^21^, prevent chitin-dependent activation of *chiS^T473A^* (**Fig. S8**). This strongly suggests that ChiS^T473A^ requires allosteric activation by liganded CBP (whereas ChiS^WT^ does not). Thus, the *chiS^T473A^* allele was used to assess allosteric, chitin-dependent activation of ChiS moving forward.

Consistent with CBP-ChiS^AF-M^ representing the liganded (active) conformation, we found that disrupting most of the intermolecular salt bridges unique to this model inhibited chitin-dependent activation (cbp^D231R^, cbp^D251R^, cbp^E401K^, chiS^T473A,R89E^, chiS^T473A,R99D^, chiS^T473A,R139E^, chiS^T473A,R146D^) (**Fig. 3, S9**). Importantly, these mutations did not affect CBP or ChiS expression (**Fig. S5**). Furthermore, flipping the charge on both sides of the interface restored chitin-dependent activation in all cases. For example, flipping the charge on either side of the CBP^D251^-ChiS^R146^ interface ablates chitin-dependent activation, while flipping the charge on both sides of the interface restores chitin-dependent activation to parent levels (**Fig. 3**). Similarly, flipping the charge on both sides of the interface restored chitin-dependent activation for CBP^D231^-ChiS^R139^, CBP^E529,E530^-ChiS^R99^, and CBP^E401^-ChiS^R89^ (**Fig. S9**). Importantly, the ChiS mutations assessed are not simply gain-of-function alleles (**Fig. S10**), indicating that this restoration is indicative of a direct interaction between the targeted residues.

**Fig. 3.**
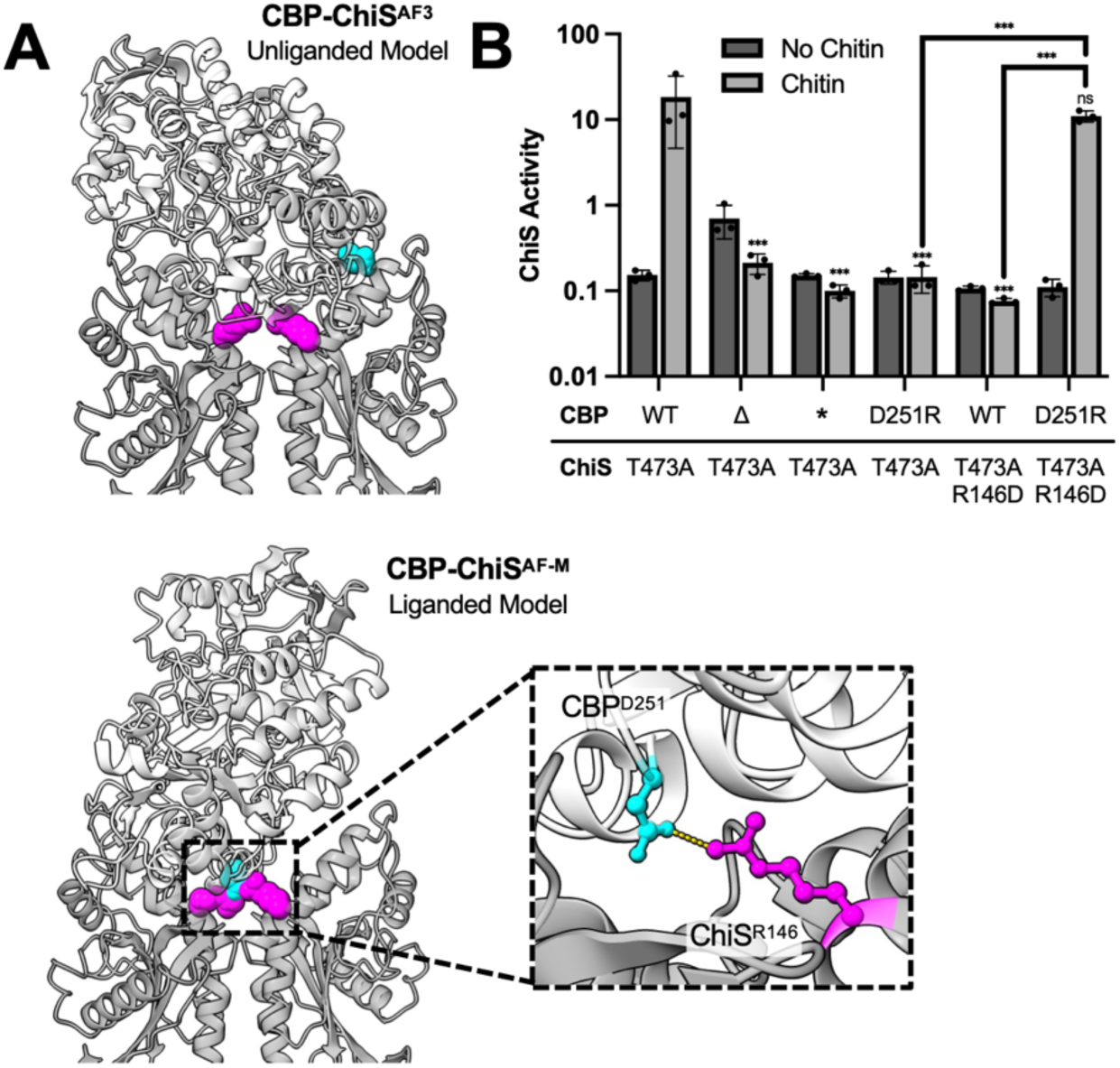
An intermolecular salt bridge specific to CBP-ChiS^AF-M^ (liganded) is necessary for allosteric activation of ChiS in the presence of chitin. (**A**) The positions of CBP^D251^ (cyan) and ChiS^R146^ (magenta) are highlighted in the CBP-ChiS^AF3^ (unliganded) and CBP-ChiS^AF-M^ (liganded) models to show that these residues only form an intermolecular salt bridge (yellow dashed line) in the latter. CBP is shown in light grey, while ChiS is shown in dark grey. (**B**) Transcriptional reporter assays to assess ChiS activity in strains harboring the indicated mutations. All strains harbored P_chb_-cbp-mCherry and P_const2_-mTFP1 reporter constructs. CBP* = CBP^W389A^ ^W539A^, a mutant allele that cannot bind to chitin. Cells were grown in either rich medium (No chitin; dark grey bars) or on chitin flakes (chitin; light grey bars) and fluorescence was determined by microscopy. Data are from three independent biological replicates (n = 300 cells analyzed per replicate) and shown as the mean ± SD. Statistical comparisons were made by one-way ANOVA with Tukey’s multiple comparison test on the log-transformed data. ns = not significant, *** = p < 0.001. Statistical identifiers directly above bars represent comparisons to the parent from the same experimental condition.

Together, these results suggest that interactions specific to CBP-ChiS^AF-M^ are dispensable for CBP-dependent allosteric repression of ChiS when chitin is absent, but necessary for allosteric activation of ChiS in the presence of chitin. This genetic analysis helps validate the hypothesis that CBP-ChiS^AF-M^ represents the conformation where ChiS is activated by chitin-bound CBP.

### Residue partner-switching facilitates CBP-dependent allosteric dual-regulation of ChiS

Thus far, we provide strong evidence for the residue-level accuracy of both the CBP-ChiS^AF3^ (unliganded) and CBP-ChiS^AF-M^ (liganded) models (**Fig. 2**, **Fig. 3**). Since ChiS activity is induced when cells are exposed to chitin, the CBP-ChiS complex must transition from the unliganded conformation to the liganded conformation when CBP binds chitin (**Movie S1**). Like many SBPs, CBP undergoes a large conformational change when binding to its chitin ligand^22,23^, which may facilitate this transition. CBP^E401^ participates at the interface of both the unliganded and liganded conformations (**Fig. 2, Fig. S9**) but maintains unique interacting partners in each model: ChiS^R141,K145^ in CBP-ChiS^AF3^ (unliganded) (**Fig. 2, 4A**) and ChiS^R89^ in CBP-ChiS^AF-M^ (**Fig. S9, 4A**). Thus, CBP^E401^ must “partner-switch”, which we hypothesize is critical for the transition between unliganded (repressed) and liganded (active) conformations.

If CBP^E401^ undergoes partner-switching, then favoring a salt-bridge interaction in the unliganded (repressed) conformation should prevent the complex from transitioning to the liganded (active) state. To test this, we revisited the charge-flip mutants for CBP^E401^-ChiS^R141,K145^. Flipping the charge on either side of the repression interface (*cbp^E401R^* or *chiS^R141E,K145E^*) only discourages the unliganded conformation (**Fig. 2B, 4B**). Accordingly, these mutations do not prevent chitin-dependent activation (**Fig. 4B**). However, flipping the charge on both sides of the interface (*cbp^E401R^ chiS^T473A,R141E,K145E^*) should restore the interaction in the unliganded conformation (*i.e.*, restoring CBP^E401^-ChiS^R141,K145^) while simultaneously discouraging the interaction in the liganded conformation (*i.e.*, disrupting CBP^E401^-ChiS^R89^). This should skew the complex towards the unliganded (repressed) conformation by preventing the partner-switch mediated transition to the liganded (active) conformation. Indeed, chitin-dependent activation is abolished in *cbp^E401R^ chiS^T473A,R141E,K145E^*(**Fig. 4B**). These results further validate the CBP-ChiS^AF3^ (unliganded) and CBP-ChiS^AF-M^ (liganded) models and suggest that residue partner-switching is a critical feature of the conformational transition associated with chitin-dependent allosteric activation.

**Fig. 4.**
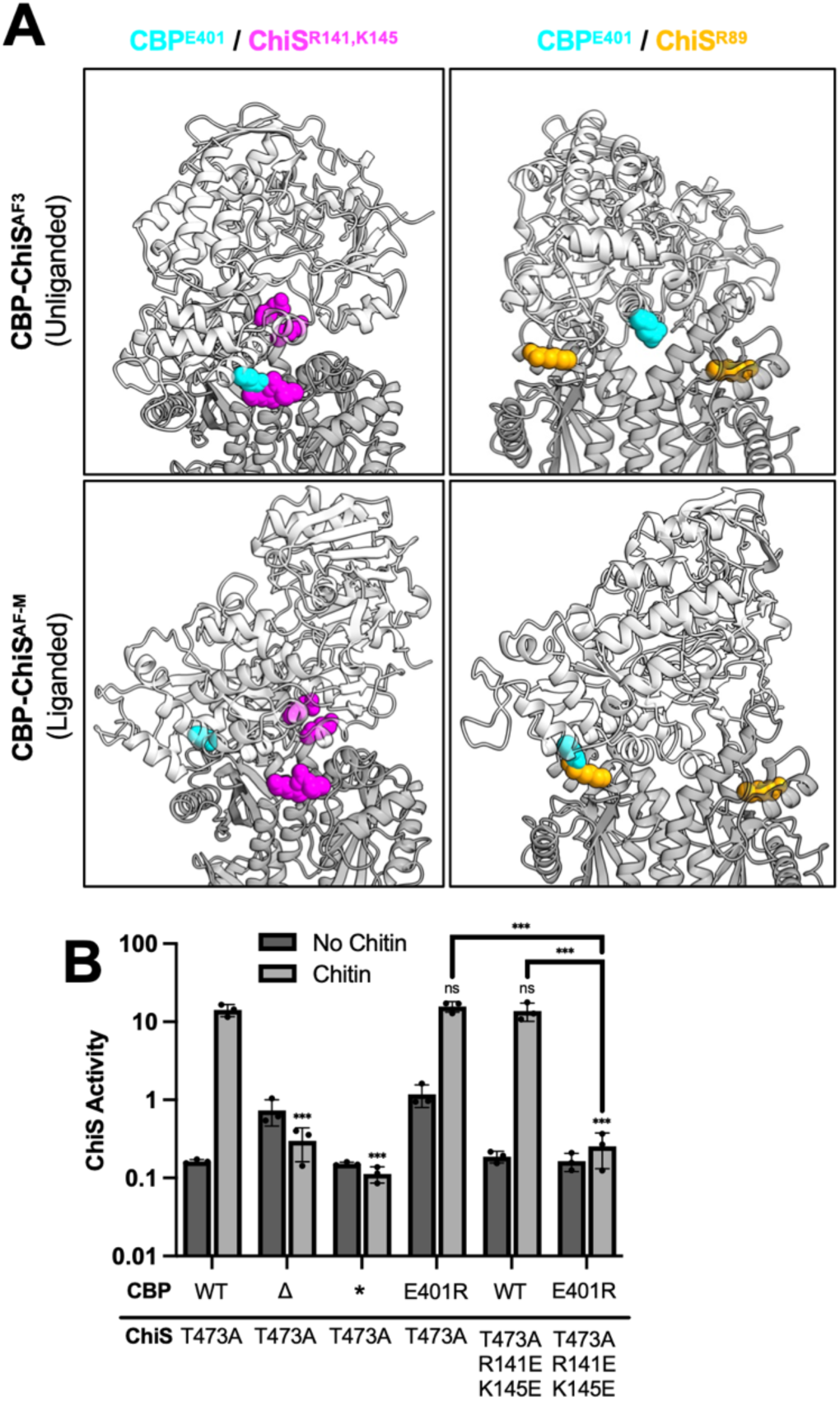
Residue partner-switching facilitates CBP-dependent allosteric dual-regulation of ChiS. CBP^E401^ undergoes partner switching to form distinct intermolecular salt bridges when transitioning from the unliganded (repressed) to liganded (active) conformations. (**A**) Models highlight that CBP^E401^ (cyan) interacts with ChiS^R141,K145^ (magenta) in CBP-ChiS^AF3^ (unliganded) and ChiS^R89^ (orange) in CBP-ChiS^AF-M^ (liganded). CBP is shown in light grey, while ChiS is shown in dark grey. (**B**) Transcriptional reporter assays to assess ChiS activity in strains harboring the indicated mutations. All strains harbored P*_chb_-cbp-mCherry* and P*_const2_-mTFP1* reporter constructs. CBP* = CBP^W389A,W539A^, a mutant allele that cannot bind to chitin. Cells were grown in either rich medium (No chitin; dark grey bars) or on chitin flakes (chitin; light grey bars) and fluorescence was determined by microscopy. Data are from three independent biological replicates (*n* = 300 cells analyzed per replicate) and shown as the mean ± SD. Statistical comparisons were made by one-way ANOVA with Tukey’s multiple comparison test on the log-transformed data. ns = not significant, *** = *p* < 0.001. Statistical identifiers directly above bars represent comparisons to the parent from the same experimental condition.

### Transitioning from the unliganded to liganded conformation is critical for ChiS-induced behaviors

ChiS activates both the chitin utilization program^6,13,15^ and a small RNA required for horizontal gene transfer by natural transformation^24^. Our results suggest that in the presence of chitin, CBP-ChiS transitions from the unliganded conformation to the liganded conformation to induce ChiS activity. To test whether this transition is required for the ChiS-regulated behaviors, we assessed our mutants for growth on insoluble chitin and for chitin-induced natural transformation. In particular, we tested the phenotype of charge flip mutations at both the liganded interface, CBP^D251^-ChiS^R146^ (**Fig. 3**), and the unliganded interface, CBP^E401^-ChiS^R141,K145^ (**Fig. 2**). We found that the ChiS-dependent activity of these mutants directly correlated with chitin growth (**Fig. 5A**) and natural transformation (**Fig. 5B**).

**Fig. 5.**
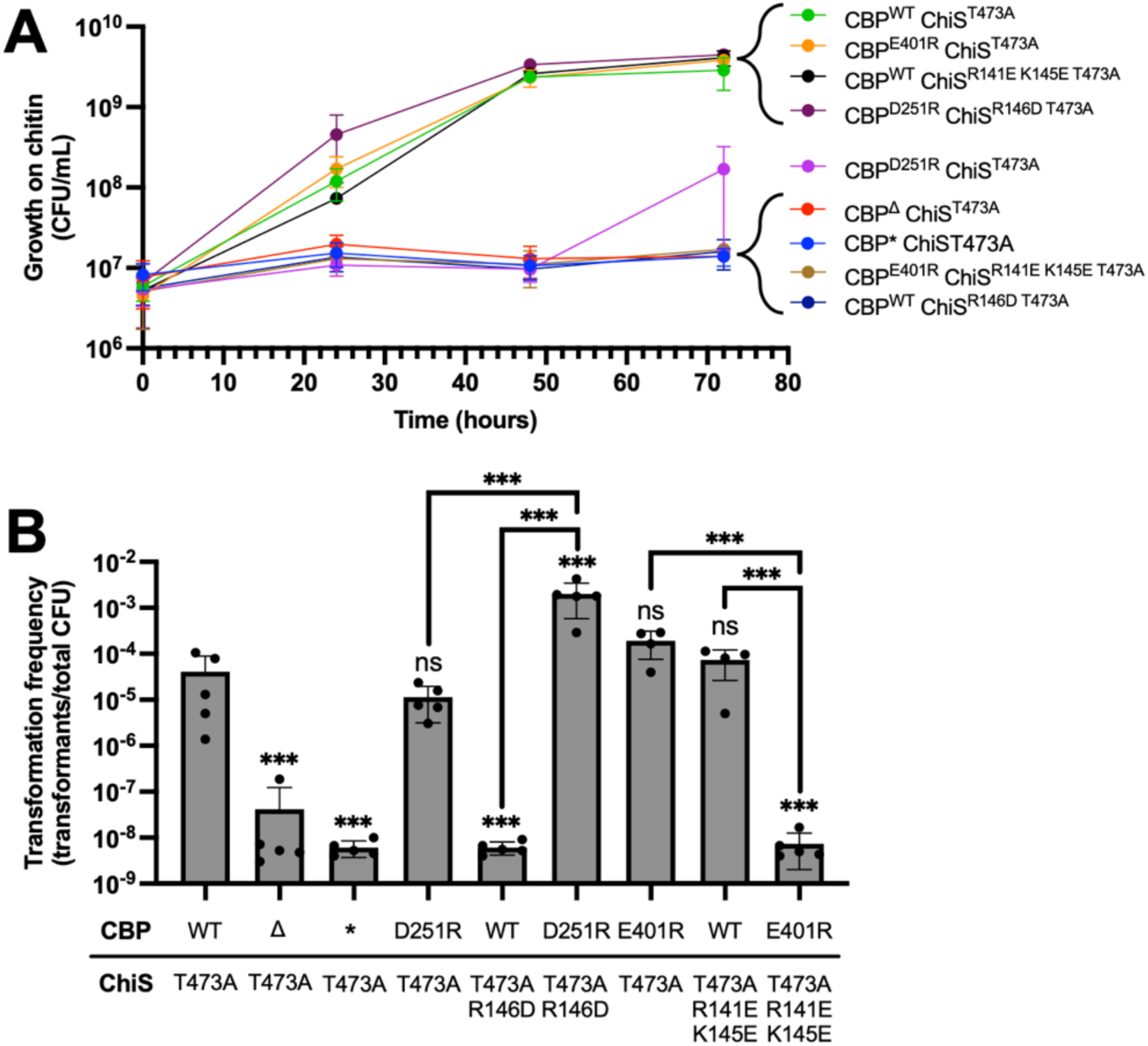
Transitioning from the unliganded (repressed) to liganded (active) conformation is critical for ChiS-induced behaviors. (**A**) The indicated strains were grown in M9 minimal media supplemented with insoluble chitin as the sole carbon source. Growth was assessed by quantitative culture at the indicated timepoints. (**B**) Chitin-induced natural transformation assays. CBP* = CBP^W389A,W539A^, a mutant allele that cannot bind to chitin. All data are from at least three independent biological replicates and shown as the mean ± SD. Statistical comparisons were made by one-way ANOVA with Tukey’s multiple comparison test on the log-transformed data. ns = not significant, *** = *p* < 0.001. Statistical identifiers directly above bars represent comparisons to the parent from the same experimental condition.

### Restrained modeling reinforces a CBP-ChiS 1:2 stoichiometry

All previously structurally characterized SBP-HK complexes adopt a 2:2 stoichiometry, where each monomer of the HK is bound by a distinct SBP^9–12^. As discussed above, we initially ruled out a 2:2 stoichiometry for CBP-ChiS based on the lower model confidence and abundance of steric clashes compared to 1:2 models (**Fig. S1**). To more rigorously test whether CBP-ChiS adopts a 1:2 or 2:2 stoichiometry, we used Chai-1^25^ to perform restrained modeling by leveraging the interface contacts that we have experimentally validated. Specifically, we generated 1:2 and 2:2 models where unliganded interface specific (CBP^E401^-ChiS^R141,K145^) or liganded interface specific (CBP^E401^-ChiS^R89^) contacts were provided as restraints (**Fig. S11A-B**). For the CBP^E401^-ChiS^R141,K145^ restrained models, the CBP^E401^-ChiS^R141,K145^ contact is maintained in both the 1:2 and 2:2 models, however, the 2:2 model contains a large number of steric clashes (**Fig. S11A**). This may suggest that steric constraints prevent CBP-ChiS from adopting a 2:2 stoichiometry in the unliganded conformation. For the CBP^E401^-ChiS^R89^ restrained models, the 2:2 model also contains a large number of steric clashes compared to the 1:2 model (**Fig. S11B**). But perhaps most notably, the CBP^E529,E530^-ChiS^R99^ interaction that we show is critical for the liganded conformation (**Fig. S9**) is not maintained in the 2:2 model, likely due to steric constraints (**Fig. S11B**). Thus, a 2:2 stoichiometry for the liganded conformation is not compatible with the empirical evidence. Altogether, this additional analysis further reinforces that CBP-ChiS likely adopts a 1:2 stoichiometry.

### Disulfide locking CBP-ChiS into the liganded (active) conformation induces ChiS activity in the absence of chitin

Thus far, our data supports the hypothesis that CBP and ChiS form a dynamic complex, that transitions between the unliganded (repressed) and liganded (active) conformations (**Movie S1**). Above, we show that skewing CBP-ChiS interactions in favor of the unliganded conformation likely prevents the structural transition associated with chitin-dependent activation (**Fig. 4**). Conversely, we hypothesized that if we could artificially stabilize CBP-ChiS interactions in the liganded conformation, we could induce ChiS activity in the absence of CBP’s chitin ligand. To that end, we introduced cysteines in CBP and ChiS at positions that are closely interacting in CBP-ChiS^AF-M^ (liganded) to “lock” the complex into the liganded (active) conformation via a disulfide bond. Specifically, we introduced cysteines at CBP^D310^ and ChiS^Q243^, whose α-carbons are <10 Å apart in CBP-ChiS^AF-M^, but >20 Å apart in CBP-ChiS^AF3^ (**Dataset 1, Fig. 6A**). As a negative control, CBP^D310C^ was paired with ChiS^Y245C^, a position that is not predicted to interact with CBP^D310^ in either the unliganded or liganded conformations (**Fig. 6A**).

**Fig. 6.**
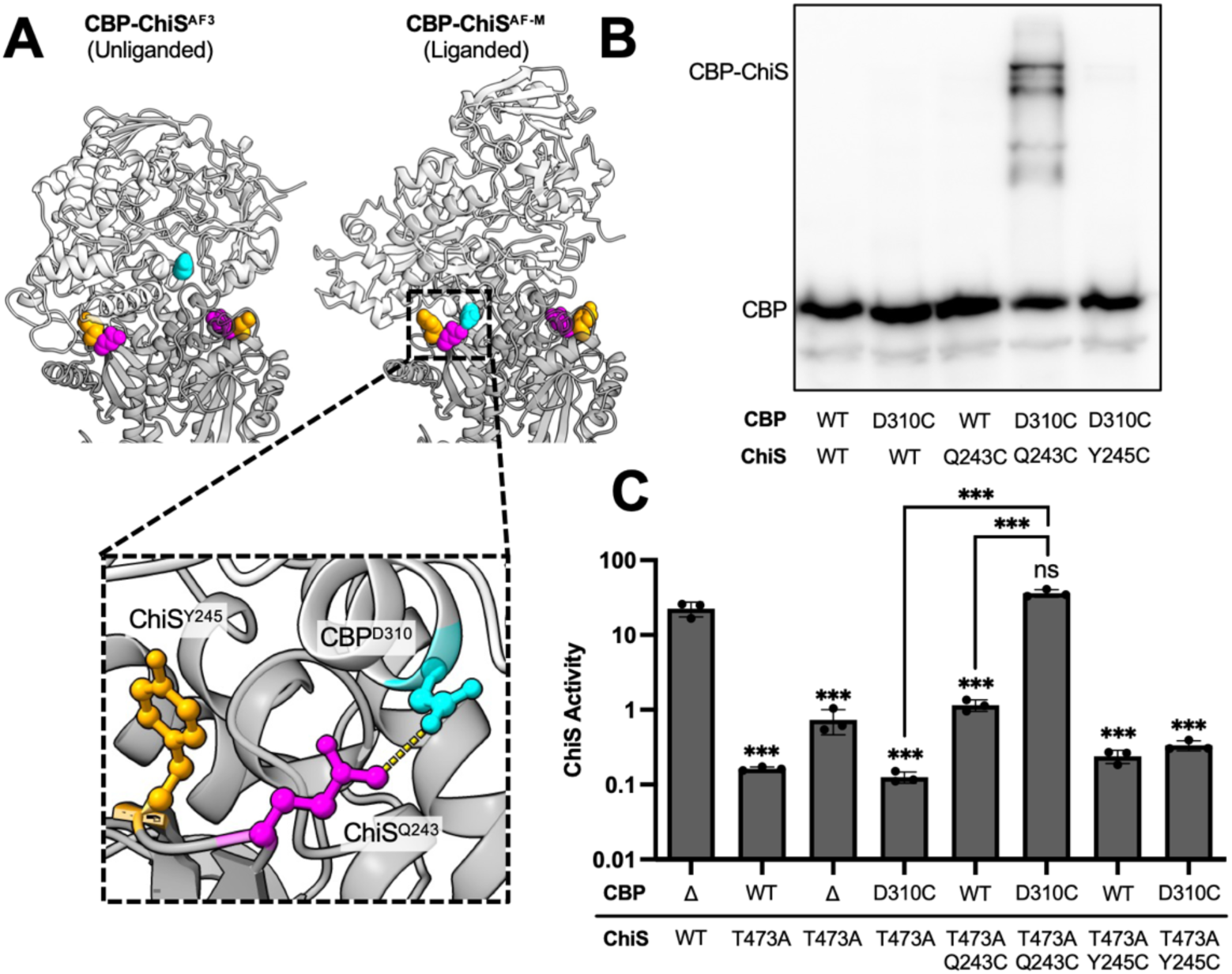
Disulfide locking CBP-ChiS into the liganded (active) conformation induces ChiS activity in the absence of chitin. (**A**) Models highlight the positions of CBP^D310^ (cyan), ChiS^Q243^ (magenta), and ChiS^Y245^ (orange) in the CBP-ChiS^AF3^ (unliganded) and CBP-ChiS^AF-M^ (liganded) models. The Inset shows that CBP^D310^ is predicted to interact with ChiS^Q243^ (yellow dashed line), but not ChiS^Y245C^. CBP is shown in light grey, while ChiS is shown in dark grey. (**B**) Non-reducing Western blot to assess disulfide crosslinking between CBP-mCherry and ChiS-FLAG using an α-mCherry primary antibody. All strains harbor *P_tac_-chiS^ΔDBD^-tetR* and *P_chb_-cbp-mCherry* constructs with the indicated mutations. (**C**) Transcriptional reporter assays to assess ChiS activity in strains harboring the indicated mutations. All strains harbored P*_chb_-cbp-mCherry* and P*_const2_-mTFP1* reporter constructs. Cells were grown in rich medium (*i.e.*, in the absence of chitin) and fluorescence was determined by microscopy. The data for the CBP^Δ^ ChiS^WT^, CBP^WT^ ChiS^T473A^, and CBP^Δ^ ChiS^T473A^ controls were duplicated from Fig. 2B and 4B and included here for ease of comparison. Data are from three independent biological replicates (*n* = 300 cells analyzed per replicate) and shown as the mean ± SD. Statistical comparisons were made by one-way ANOVA with Tukey’s multiple comparison test on the log-transformed data. ns = not significant, *** = *p* < 0.001. Statistical identifiers directly above bars represent comparisons to the first sample (CBP^Δ^ ChiS^WT^).

First, disulfide crosslinking was assessed by non-reducing Western blots. When cells harbor only a single cysteine knock-in (CBP^D310C^ or ChiS^Q243C^), disulfide crosslinking is not observed (**Fig. 6B, S12A**), despite robust CBP-ChiS interactions in these strains, as evidenced by NHS-ester crosslinking with disuccinimidyl suberate (DSS) (**Fig. S12B**). However, in cells harboring both CBP^D310C^ and ChiS^Q243C^, robust disulfide crosslinking is observed (**Fig. 6B, S12A**). Importantly, disulfide crosslinks are not observed in the CBP^D310C^ ChiS^Y245C^ negative control (**Fig. 6B, S12A**), despite robust CBP-ChiS interactions (**Fig. S12B**). Importantly, CBP^D310C^-ChiS^Q243C^ disulfide crosslinking occurs in cells grown in rich medium (*i.e.*, in the absence of chitin). This suggests that CBP-ChiS dynamically samples the active conformation even in the absence of its chitin ligand, and that disulfide crosslinking of CBP^D310C^-ChiS^Q243C^ locks the complex into the liganded conformation.

Next, we tested whether the liganded conformation is sufficient for ChiS activity. We found that *cbp^D310C^ chiS^T473A,Q243C^* exhibited strong ChiS activity, even in the absence of chitin (**Fig. 6C**). Conversely, this was not observed for strains harboring only a single cysteine (*cbp^D310C^* or *chiS^T473A,Q243C^*) or the non-disulfide crosslinked cysteine pair (*cbp^D310C^-chiS^T473A,Y245C^*) (**Fig. 6C**). Importantly, the *chiS^Q243C^* and *chiS^Y245C^* cysteine mutations do not diminish ChiS activity (**Fig. S7**). Together, these results strongly suggest that stabilizing the CBP-ChiS^AF-M^ (liganded) conformation, even in the absence of chitin, is sufficient to induce ChiS activity. Moreover, they suggest that the CBP-ChiS complex dynamically samples both the unliganded and liganded conformations, with the latter likely being stabilized when CBP binds to its chitin ligand.

### AF-Mv3 predicts distinct conformations for other SBP-HK signaling complexes

Our results suggest that AlphaFold can reveal the distinct conformations associated with dual allosteric regulation of CBP-ChiS (**Fig. 1**). However, we required two separate modeling algorithms (AF3 and AF-Mv3) to reveal these distinct conformations. We hypothesized that AF3 and AF-Mv3 might each be able to model both conformations, but algorithm-specific criteria may prioritize one conformation to effectively hide the other conformation from the top-ranked models. We hypothesized that analyzing the lower ranked models or changing the modeling parameters may, however, allow us to uncover both conformations using each algorithm.

For AF3, we found that analyzing the lower ranked models did not reveal the alternative (liganded) conformation. However, we found that AF3 modeling the disulfide lock residues, CBP^D310C^-ChiS^Q243C^, switches the output model from the unliganded conformation to the liganded conformation that was initially revealed by AF-Mv3 (**Fig. S13A,C**). Importantly, AF3 continues to predict the unliganded conformation for CBP^D310C^-ChiS^Y245C^ (**Fig. S13B-C**), which we empirically demonstrate cannot form a disulfide crosslink (**Fig. 6**). While this modeling supports that AF3 can predict the liganded conformation, it cannot do so agnostically and requires prior knowledge of interfacial contact residues to reveal this originally “hidden” conformation.

In contrast to AF3, further analysis of the models obtained by AF-Mv3 revealed that both conformations of CBP-ChiS were present when all of the ranked models were analyzed (**Fig. S14**). AF-Mv3’s ability to agnostically predict both CBP-ChiS conformations suggested that it might be able to uncover multiple conformations for other protein complexes *in silico*. Specifically, we sought to determine whether AF-Mv3 could reveal the allosteric activation mechanism for other SBP-HK signaling complexes that remain structurally uncharacterized^4^. We found that AF-Mv3 predicted two distinct conformations for *Agrobacterium tumefaciens* ChvE-VirA^26^ and *Bordetella pertussis* BctC-BctE^8^ (**Fig. S14**). Like CBP-ChiS, the orientation of the SBP changes with respect to the HK to produce two distinct conformations for ChvE-VirA and BctC-BctE (**Fig. S14, S15A, Movie S2-S3**). Thus, it is tempting to speculate that these distinct conformations represent the unliganded and liganded states of these signaling complexes.

While the molecular mechanism of BctC-BctE signal transduction remains poorly understood, ChvE-VirA is relatively well-characterized. In particular, one study has defined a number of ChvE mutant alleles that alter allosteric activation of VirA^5^. Specifically, it uncovered ChvE mutations that either (1) prevented VirA activation in the presence of ligand or (2) induced VirA activity in the absence of ligand. We hypothesize that the former represents residues required for the liganded interface, while the latter represent residues that stabilize the unliganded interface (and thus, their mutation skews the complex into the liganded/active conformation). Consistent with this, these residues lie at the ChvE-VirA interface in the two conformations predicted by AF-Mv3 (**Fig. S15B**). The residues required for ChvE-VirA activation in the presence of ligand are closer to the interface of one model, while the residues that, when mutated, induce ChvE-VirA activation in the absence of ligand are closer to the interface of the other model (**Fig. S15C**, **Movie S2**), which supports their opposing effects on ChvE-VirA activity. Furthermore, analyzing both ChvE-VirA conformations for predicted contacts reveals potential evidence for residue partner-switching (**Fig. S15D**) akin to what is observed for CBP-ChiS (**Fig. 4**). Importantly, mutation of the ChvE residues involved in this partner-switching have opposing phenotypic effects, where mutation of one residue, ChvE^L181^, prevents activation in the presence of ligand, while mutation of the other, ChvE^A186^, induces VirA activity in the absence of ligand^5^. Altogether, this analysis provides compelling evidence for the biological relevance of the two ChvE-VirA conformations predicted by AF-Mv3. More broadly, these results highlight how AF-M modeling can help reveal the molecular basis for allosteric signal transduction.

## DISCUSSION

Through a combination of structural modeling and complementary genetic and biochemical approaches, this study has uncovered the molecular mechanism of the dual allosteric switch regulating the chitin utilization program of *V. cholerae*. Our results indicate that the CBP-ChiS complex undergoes a large conformational change upon ligand-binding to induce the activity of the ChiS sensor kinase. More broadly, this work highlights how structural modeling combined with structure-function analysis can rapidly reveal the molecular basis for signal transduction.

SBP-HK complexes facilitate signal transduction to a variety of environmental signals^4^. In these systems, liganded SBPs must induce a conformational change in their cognate HK to induce their activity. The molecular basis for this activity, however, varies widely^9–12^, highlighting the lack of a broadly conserved allosteric activation mechanism. All currently characterized SBP-HK systems adopt a 2:2 heterotetrameric complex in the liganded form. By contrast, our data suggests that liganded CBP-ChiS uniquely adopts a 1:2 stoichiometry. While a 1:2 stoichiometry has not been observed among other SBP-HKs, a similar arrangement is predicted for maltose binding protein (MBP)-dependent activation of the Tar chemoreceptor^27^. Furthermore, SBPs typically function as monomers to transport ligands in association with their cognate ABC transporter complex^28–30^.

For all structurally characterized SBP-HK systems, the orientation of the SBP with respect to the HK remains relatively unchanged upon ligand binding. This includes LuxP-LuxQ^9^, TorT-TorS^10^, XylFII-LytS^12^, and HptA-HptS^11^. By contrast, we show that CBP-ChiS undergo an interface switch upon ligand-binding, where the orientation of CBP exhibits a large rotation relative to the ChiS dimer (**Movie S1**). Structural modeling suggests that similar interface switches also facilitate the activation of ChvE-VirA and BctC-BctE (**Movie S2-S3**). Thus, interface switching may represent a novel mechanism for allosteric activation of signal transduction in SBP-HKs.

Our results suggest that structural modeling can rapidly reveal the allosteric activation mechanism of protein complexes. In particular, we show that AF-Mv3 can agnostically model distinct conformations for a number of SBP-HKs. This was not observed when using earlier versions of AF-M (v1 or v2) or AF3, which generated CBP-ChiS models in only one conformation. Furthermore, AF3 yielded low confidence models for ChvE-VirA, suggesting that AF-Mv3 may be better at modeling some protein complexes. AF3, however, was able to model CBP in its unliganded state (**Fig. S3**) and reveal which CBP-ChiS complex represents the ChiS-repressed conformation. This would not have been possible with AF-Mv3 alone. While AF-Mv3 revealed multiple conformations for CBP-ChiS, the CBPs in these models were identical and matched the crystal structure of liganded CBP. So, employing multiple structural modeling algorithms (*e.g.*, AF-M^18^, AF3^19^, Chai-1^25^, RosettaFold2^31^, and RosettaFold All Atom^32^) and comparing between models may help reveal allosteric conformations in other protein complexes. Furthermore, recent work highlights that clustering and/or subsampling of multiple sequence alignments (MSAs) can help elucidate protein conformational landscapes^33–36^, which may help accelerate the discovery of additional allosteric activation mechanisms moving forward.

## METHODS

### Structural modeling with AlphaFold

The initial CBP-ChiS models (**Fig. 1**) were generated using the AF-Mv3 algorithm^18^ in ColabFold^16^ on Indiana University’s BigRed200 and Quartz Supercomputers or using the AlphaFold3 algorithm^19^ on the AlphaFold webserver. For AF-Mv3, multiple sequence alignments (MSAs) were generated by MMseqs2^37^ and five relaxed models were generated from a single seed using 50 recycles with tolerance set to 0.2. The 5 top ranked models from 5 independent runs were used for subsequent analysis. To identify distinct protein conformations using AF-Mv3 (**Fig. S14** and **S15**) each query generated 1 unrelaxed model for 25 independent random seeds using 30 recycles with tolerance set to 1.0.

Models were analyzed using ChimeraX^38^ (v1.9), and contacts were determined using the “contacts” and “h-bonds” tools. Parameters for “contacts” included a VDW overlap of ≥ -0.4 Å and intramodel interactions only (intermodel, intramolecule, intraresidue contacts were excluded). Parameters for “h-bonds” included a distance tolerance of 5 Å, angle tolerance of 20°, and salt-bridges only. For a complete list of predicted electrostatic contacts, see **Dataset S1**. Structural models are available at ModelArchive^39^ (Accession: ma-dal-cbpchis).

### Bacterial strains and culture conditions

Bacterial strains were grown in LB Miller broth at 30°C rolling and on LB Miller agar. When necessary, the medium was supplemented with spectinomycin (200 μg/ml), kanamycin (50 μg/mL), trimethoprim at 10 (μg/mL), chloramphenicol (1 μg/mL), carbenicillin (100 μg/mL), erythromycin (10 μg/mL), sulfamethoxazole (100 μg/mL), tetracycline (0.5 μg/mL), and/or zeocin (100 μg/mL) as appropriate.

All strains were derived from the *Vibrio cholerae* El Tor strain E7946. Mutant constructs were created using splicing-by-overlap-extension (SOE) PCR exactly as previously described^40^. All chromosomally integrated ectopic expression constructs were generated exactly as previously described^41^. Mutant constructs were transformed into *V. cholerae* using natural transformation. Competence was artificially induced in cells via ectopic expression of TfoX, the master regulator of competence, using either an IPTG-inducible plasmid (pMMB67EH-*tfoX*) or via a chromosomally integrated arabinose-inducible construct (P*_BAD_*-*tfoX*). The ectopic *tfoX* expression construct was either cured (for pMMB67EH-*tfoX*) or deleted (for P*_BAD_*-*tfoX*) from all strains prior to their use in experiments. All mutations were verified by PCR and/or sequencing.

For a detailed list of all strains, see **Table S1**. For a complete list of oligos used to generate mutant constructs specific to this study, see **Table S2**. The CBP^Δ^ construct used throughout the study consists of the CBP signal sequence translationally fused to mCherry (*i.e.*, P*_chb_-cbp^1-40^-mCherry)*.

### Microscopy

Cells were imaged on an inverted Nikon Ti-2 microscope with a Plan Apo 60x objective lens, the appropriate filter cubes (FITC, YFP, mCherry, and/or CFP), a Hamamatsu ORCA Flash 4.0 camera, and Nikon NIS Elements imaging software. All image analysis was performed using the MicrobeJ^42^ plugin (v5.10m) in Fiji^43^ (v1.53k).

### ChiS activity assays

Single colonies were inoculated into LB medium and incubated at 30°C for 18 hours rolling to generate overnight cultures. To assess chitin-independent ChiS activity by plate reader, cells were washed and resuspended in Instant Ocean (IO) medium (7 g/L, Aquarian Systems) to an OD_600_ of ∼5.0. Then, 200 μL of cell suspension was transferred to two wells (technical replicates) per strain in a flat-bottom 96-well plate and fluorescence was determined on a Biotek H1M plate reader. The fluorescence monochromator settings for excitation/emission were: 446/491 for mTFP1 and 580/610 for mCherry. Background fluorescence was determined by analyzing a non-fluorescent strain and was subtracted from all experimental samples. Technical replicate measurements were averaged. Then, CBP-mCherry fluorescence was normalized by mTFP1 fluorescence to determine the ChiS activity for each biological replicate.

To assess chitin-independent ChiS activity by microscopy, overnight cultures were washed and resuspended in IO to an OD_600_ of ∼1.0. Then, 2 μL of this cell suspension was placed under a 0.3% Gelzan IO pad. Cells were imaged and the average fluorescence signal per cell was determined using MicrobeJ. Background fluorescence was determined by analyzing a non-fluorescent strain and was subtracted from all experimental samples. For each cell, the CBP-mCherry fluorescence was normalized by mTFP1 fluorescence to control for intrinsic noise in gene expression. The geometric mean of 300 cells was reported for each biological replicate as the ChiS activity.

To assess chitin-dependent ChiS activity by microscopy, overnight cultures were subcultured into fresh LB and grown to an OD_600_ of ∼1.0. Cells were then washed and resuspended to an OD_600_ of 1.0 in IO. Then, 100 μL of cell suspension was mixed with 750 μL of IO and 150 μL of insoluble chitin slurry (5% w/v insoluble chitin flakes in IO). Reaction tubes were shaken at 30°C for 48 hours to allow for chitin-induction. Then, reactions were vortexed to release the cells from chitin and the supernatant was transferred to a fresh Eppendorf tube to concentrate the cells in IO. Next, cells were placed under a 0.3% Gelzan IO pad and imaged and analyzed exactly as described above for chitin-independent ChiS activity assays.

### PopZ-linked apical recruitment (POLAR) assays

Single colonies were inoculated into LB medium supplemented with 0.1 μM IPTG to induce P*_tac_*-*chiS*-*msfGFP*-*H3H4* expression. Where indicated, samples were also supplemented with 0.05% arabinose to induce P*_BAD_*-PopZ expression to induce polar relocalization of the H3H4-tagged bait. In all strains, the cell curvature determinant *crvA* was deleted because ectopic expression of ChiS induces strong cell curvature^44^, which confounds the analysis. Importantly, the *chiS*-msfGFP-H3H4 construct is nonfunctional for *P_chb_*-induction, thus, the *P_chb_*-*cbp*-*mCherry* construct is expressed at a basal and equivalent level in all strains. Cultures were incubated at 30°C for 18 hours rolling. Then, cells were subcultured the following day in fresh medium and grown for 1 hour at 30°C rolling. Cells were then washed and resuspended in IO to an OD_600_ of ∼1.0. Then, 2μL of cell suspension was placed under a 0.3% Gelzan IO pad and imaged. Fluorescence localization within cells was determined using MicrobeJ’s “Maxima” function, where foci in mCherry and FITC channels were called using the “Point” mode of detection with a tolerance of 50. Cell boundaries were determined using “Fit Shape > Rod-Shaped” segmentation and foci outside of cell boundaries were excluded. Heatmaps of fluorescence localization were generated by analyzing 300 cells per replicate. To quantify polar localization, the distance of fluorescence to the nearest cell pole was measured. Fluorescence ≤0.2 μm from the cell pole was conservatively defined as polarly localized.

### Western Blotting

To assess the impact of point mutations on CBP and ChiS expression, the native *chiS* and *cbp* genes were deleted and complemented with ectopic *P_chb_-cbp-mCherry* or *P_chiS_-chiS-FLAG* constructs. To assess CBP expression, cells were resuspended to an OD_600_ of 50 in ice cold IO. Cells were then chemically lysed by adding 10% FastBreak, lysozyme (0.5 mg/mL), and Benzonase (0.5 unit/μL) and incubating for 15 minutes at room temperature. Then, samples were mixed 1:1 with 2X reducing SDS-PAGE sample buffer (222 mM Tris HCl pH 6.8, 26 % glycerol, 3.9 % SDS, 0.022 % bromophenol blue, 0.715 M beta-mercaptoethanol, 160 mM DTT). To assess ChiS expression, cells were resuspended to an OD_600_ of 100 in ice cold IO. Samples were then mixed 1:1 with 2X reducing SDS-PAGE sample buffer and incubated at 100°C for 5 minutes.

For CBP-ChiS crosslinking (disulfde and DSS), to ensure CBP expression levels were equivalent between strains, we used a previously described ChiS construct^24^ where the native DNA-binding domain was deleted (ΔDBD) and replace by TetR. Strains were grown in LB medium supplemented with 0.04% arabinose to induce *P_bad_-chiS^ΔDBD^-tetR* expression at 30°C for 18 hours rolling. Strains were then subcultured into fresh medium and grown at 30°C to an OD_600_ of ∼4.0. Cells were then washed in ice cold IO.

For disulfide crosslinked samples, cells were resuspended to an OD_600_ of 100 in ice cold IO. Next, cells were chemically lysed by adding 10% FastBreak, lysozyme (0.5 mg/mL), and Benzonase (0.5 unit/μL) and incubating for 15 minutes at room temperature. To assess disulfide crosslinking, samples were mixed 1:1 with 2X non-reducing SDS-PAGE sample buffer (222 mM Tris HCl pH 6.8, 26 % glycerol, 3.9 % SDS, 0.022 % bromophenol blue). To reduce disulfide crosslinks prior to gel electrophoresis, samples were mixed 1:1 with 2X reducing SDS-PAGE sample buffer.

For DSS crosslinked samples, cells were resuspended in ice cold IO, mixed with 5 mM DSS, and nutated at room temperature for 30 minutes. Samples were then quenched with 45mM Tris HCl pH 7.5 and nutated at room temperature for 15 minutes. Samples were then washed and resuspended to an OD_600_ of ∼100 in ice cold IO. Next, cells were chemically lysed exactly as described above. Then, lysates were mixed 1:1 with 2X reducing SDS-PAGE sample buffer.

Next, 5-15μL of each sample was electrophoretically separated on a 7.5% SDS-PAGE gel. Proteins were then transferred to a polyvinylidene difluoride (PVDF) membrane. Membranes were blocked in TBST (20 mM Tris HCl pH 7.5, 50 mM NaCl, 0.05% Tween 20) with 5% milk powder for 1 hour rocking at room temperature. Membranes were then incubated with 1:10,000 rabbit polyclonal α-mCherry primary antibody or 1:1,000 rabbit polyclonal α-FLAG primary antibody in TBST with 2% milk powder rocking at room temperature overnight. Membranes were then washed with TBST before a 2-hour incubation with 1:10,000 α-rabbit horseradish peroxidase-conjugated secondary antibody. Membranes were then washed and incubated with Pierce ECL Western Blotting Substrate before chemiluminescence imaging on a ProteinSimple FluorChem R instrument.

### Insoluble chitin growth assays

Single colonies were inoculated into LB medium and incubated at 30°C for 18 hours rolling. Then, cells were subcultured into fresh LB and grown to an OD_600_ of ∼1.0. Cells were washed and resuspended to an OD_600_ of 1.0 in M9 minimal medium (1X M9 salts, 2 mM MgSO_4_, 0.1 mM CaCl_2_, 0.03 mM FeSO_4_). Then, 10 μL of cell suspension was added into 1 mL reactions containing 850 μL of M9 minimal medium and 150 μL of chitin slurry. Reactions were shaken at 30°C for a total of 72 hours. Growth was assessed at 0, 24, 48, and 72 hours by quantitative dilution plating.

### Chitin-induced natural transformation assays

Single colonies were inoculated into LB medium and incubated at 30°C for 18 hours rolling. Cells were then subcultured into fresh LB and grown to an OD_600_ of ∼1.0. Cells were washed and resuspended to an OD_600_ of 1.0 in IO. Then, 100 µL of cell suspension was added into 1mL reactions containing 750 μL of IO and 150 μL of chitin slurry. Reaction tubes were incubated statically at 30°C for 18 hours. Then, 550 μL of the supernatant was removed without disturbing the settled chitin. Next, 100 ng of transforming DNA (ΔVC1807::Spec^R^) was added. For each strain, control reactions were performed where no DNA was added. Reaction tubes were incubated statically at 30°C for 5 hours. Next, 500 μL of LB medium was added to each reaction, and outgrown by incubating at 37°C for 3 hours shaking. Reactions were then dilution plated on LB supplemented with spectinomycin to quantify transformants, and plain LB to quantify total viable counts. Transformation frequency is defined as the CFU/mL of transformants divided by the CFU/mL of total viable counts.

### Statistical comparisons

All statistical comparisons were made in Graphpad Prism (v10) and the statistical tests used are indicated in each figure legend. See **Dataset S2** for a comprehensive list of descriptive statistics and statistical comparisons.

## ACKNOWLEDGEMENTS

We thank Virginia Green and Catherine Klancher for constructive comments on the manuscript and helpful discussions. This work was supported by grant R35GM128674 from the National Institutes of Health to ABD.

---

**Fig. S1.**
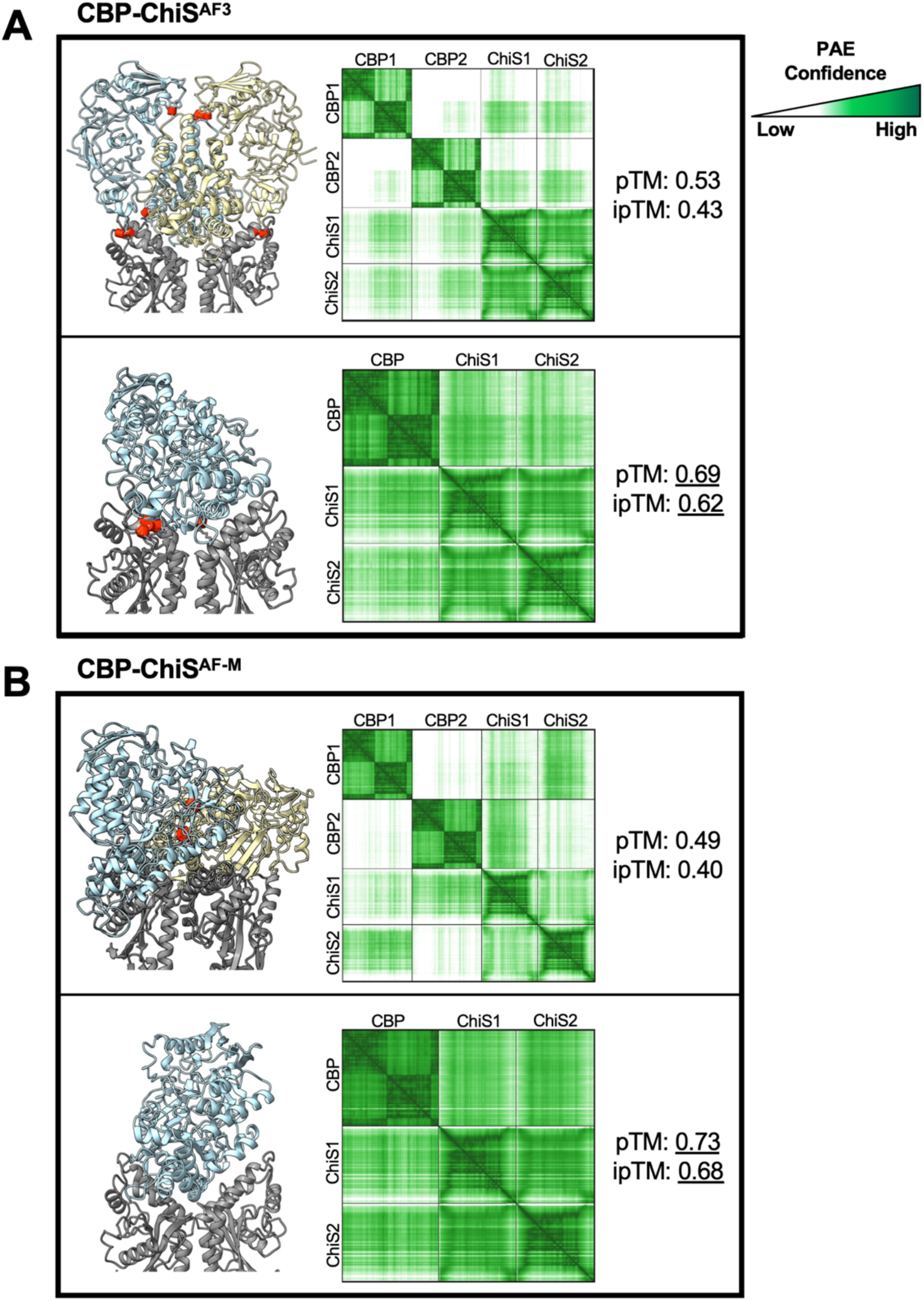
CBP-ChiS modeling at 2:2 vs 1:2 stoichiometry. Top ranked models of CBP (blue & yellow)-ChiS (grey) interactions at 2:2 (upper box) and 1:2 (lower box) stoichiometries using (**A**) AF3 and (**B**) AF-Mv3. Red cylinders on models demarcate steric clashes. <u>P</u>redicted <u>A</u>ligned <u>E</u>rror (PAE) plots are colored by confidence. Quantitative metrics (pTM and ipTM scores) for each model are shown at the right.

**Fig. S2.**
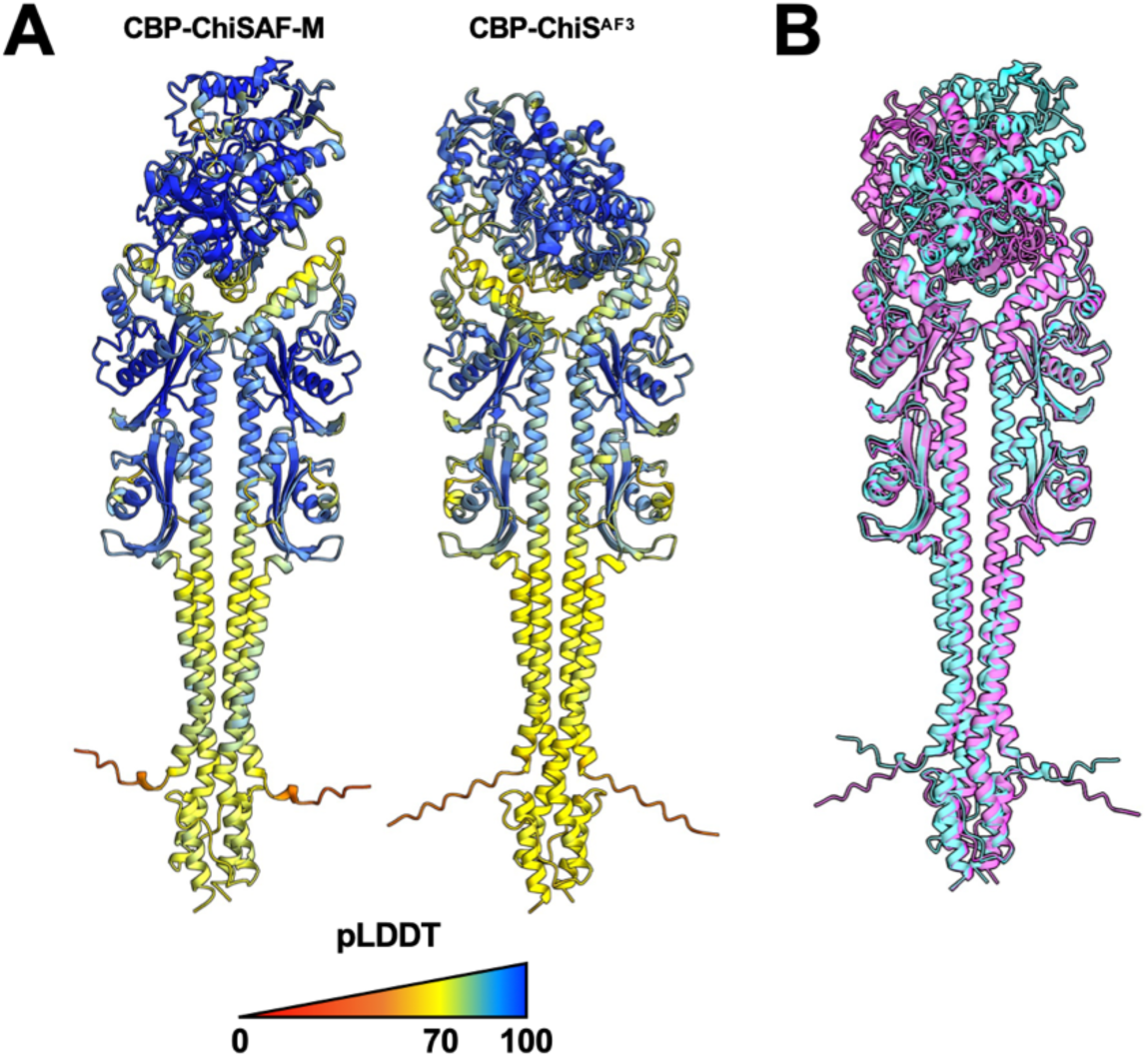
Different AlphaFold algorithms predict two distinct conformations for CBP-ChiS. (**A**) Top ranked models of CBP-ChiS^AF-M^ and CBP-ChiS^AF3^ colored by pLDDT score. CBP’s orientation is rotated relative to ChiS dimer. (**B**) Overlay of CBP-ChiS^AF-M^ and CBP-ChiS^AF3^ models to highlight the distinct orientation of CBP within each model. The CBP-ChiS^AF-M^ model is colored in cyan, while the CBP-ChiS^AF3^ model is colored in magenta.

**Fig. S3.**
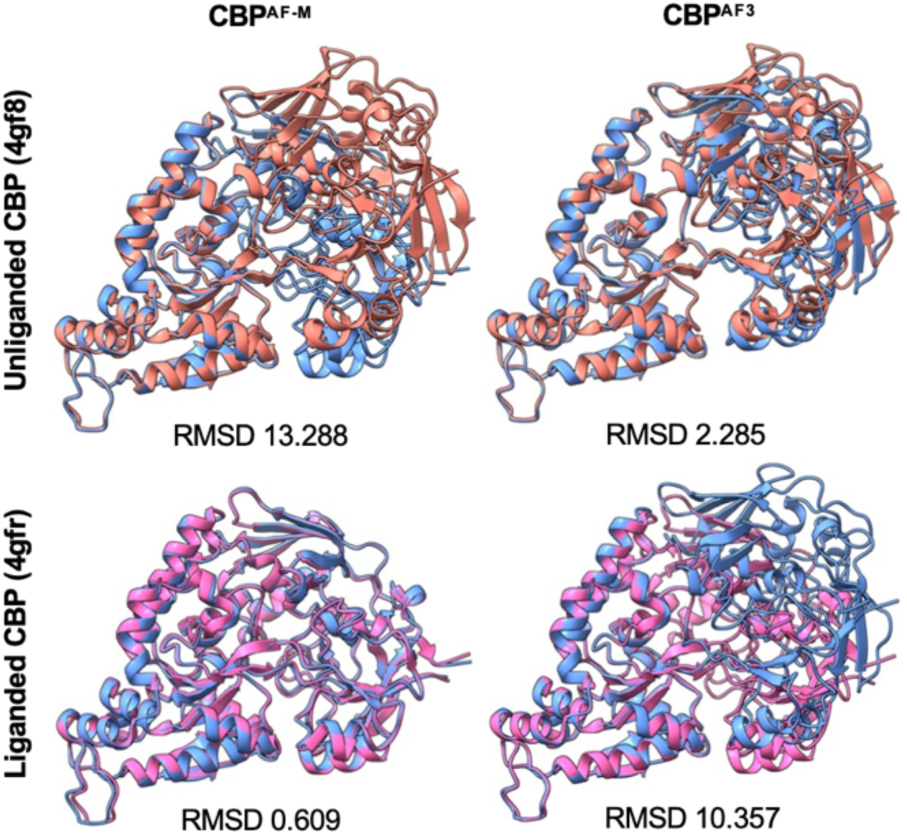
Alignment of CBP from CBP-ChiS^AF3^ and CBP-ChiS^AF-M^ to solved crystal structures reveals the liganded state of each AlphaFold model. CBP extracted from either the CBP-ChiS^AF-M^ or CBP-ChiS^AF3^ model (blue) were aligned to either unliganded CBP (4gf8, salmon) or liganded CBP (4gfr, magenta) as indicated. Untrimmed RMSD scores are given for each alignment.

**Fig. S4.**
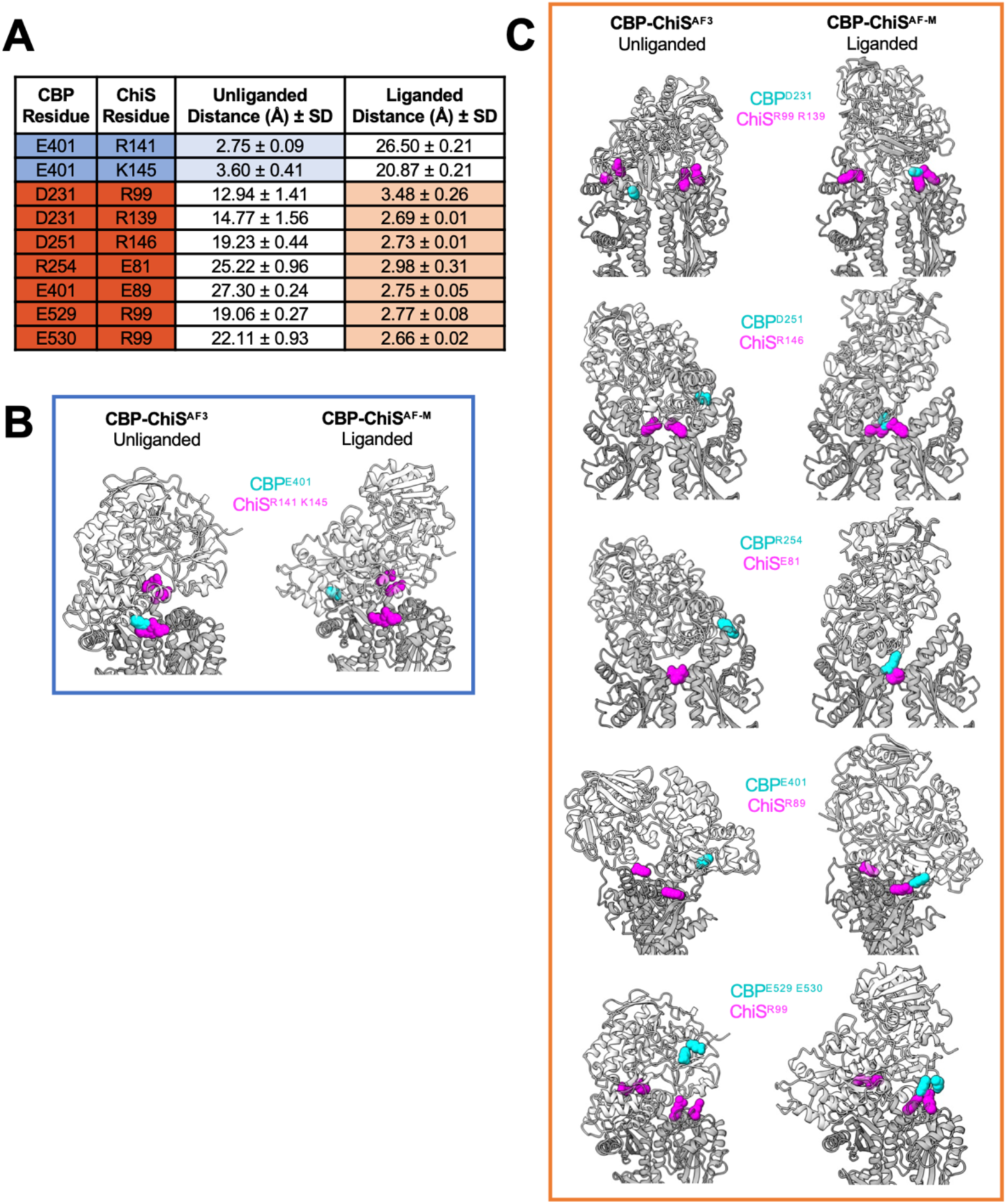
Defining putative intermolecular salt bridges unique to each AlphaFold model. The top models from five independent CBP-ChiS^AF-M^ and CBP-ChiS^AF3^ runs were analyzed to define putative intermolecular salt bridges unique to each model. (**A**) The average of the smallest distance between side chain atoms of the indicated residue pairs are shown ± SD. Residue pairs unique to (**B**) CBP-ChiS^AF3^ and (**C**) CBP-ChiS^AF-M^ residue pairs are mapped onto each AlphaFold model. CBP is colored light grey (indicated residues in cyan), while ChiS is colored dark grey (indicated residues in magenta).

**Fig. S5.**
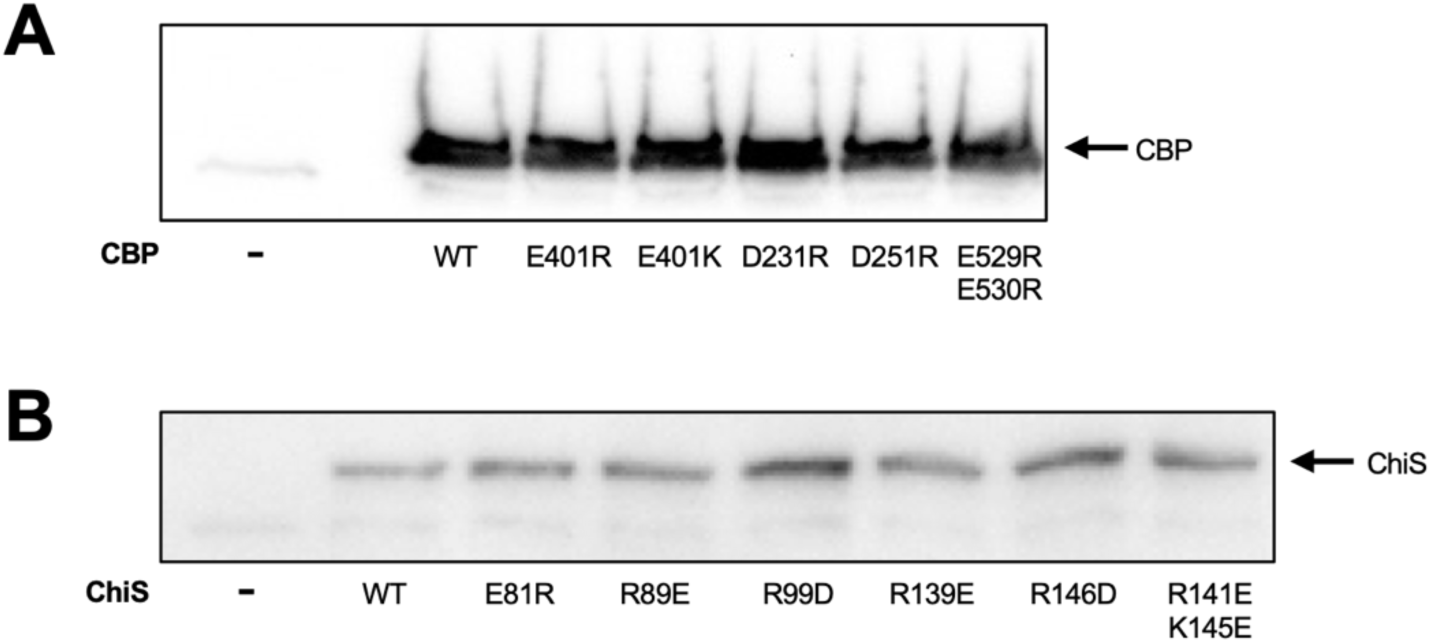
Point mutations to CBP and ChiS do not affect their expression level. Reducing western blots were used to assess protein levels of the indicated CBP and ChiS mutants. Data are representative of two independent biological replicates. “-“ demarcates a control lysate that lacks a tagged CBP/ChiS allele. (**A**) Strains harbor ectopic *P_chb_-cbp-mCherry* constructs containing the indicated CBP mutation and the native *chiS* and *cbp* genes deleted. Western blots were probed with α-mCherry primary antibody. (**B**) All strains harbor ectopic *P_chiS_-chiS-1X FLAG* constructs containing the indicated ChiS mutation and the native *chiS* and *cbp* genes deleted. Western blots were probed with α-FLAG primary antibody.

**Fig. S6.**
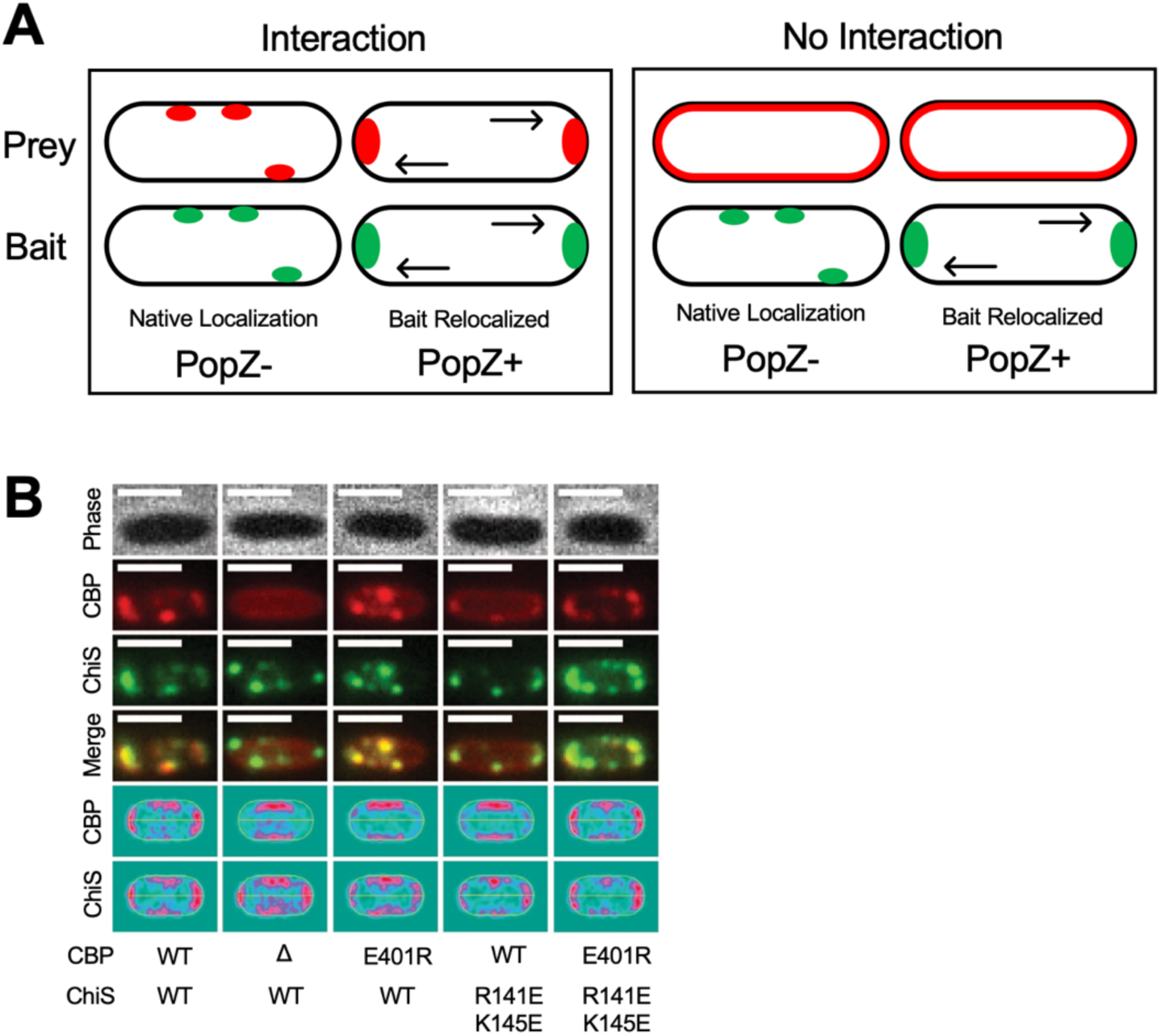
A salt bridge interaction specific to the CBP-ChiS^AF3^ (unliganded) model is necessary for allosteric repression of ChiS in the absence of chitin. (**A**) Schematic representation of the POLAR assay. (**B**) Native CBP-ChiS interactions were assessed by fluorescence microscopy, using cells without PopZ induction. Cells contained cbp-mCherry and chiS-msfGFP-H3H4 constructs with the indicated mutations as well as an inducible PopZ construct (P_BAD_-popZ). CBP^Δ^ = CBP signal sequence translationally fused to mCherry (P*_chb_-cbp^1-40^-mCherry)*. Representative images are shown. Scale bars, 2 µm. Heat maps represent the localization of mCherry and GFP fluorescence in 300 cells.

**Fig. S7.**
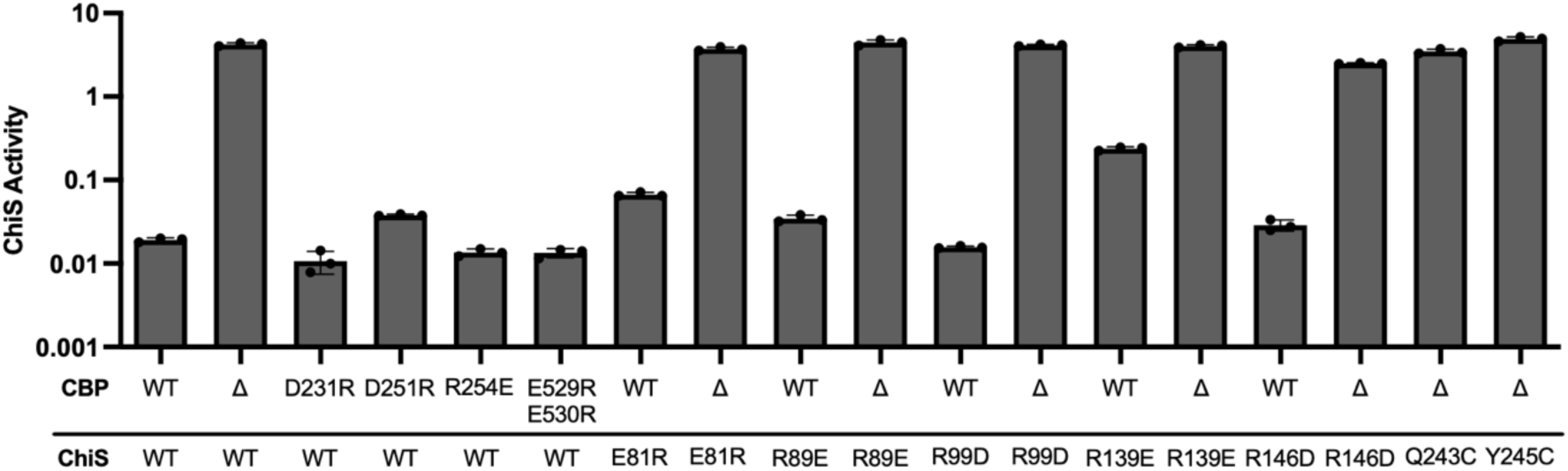
Mutating residues specific to the CBP-ChiS^AF-M^ (liganded) model do not affect CBP-dependent repression of ChiS in the absence of chitin. Transcriptional reporter assays to assess ChiS activity in strains harboring the indicated mutations that specifically disrupt the CBP-ChiS interface in the CBP-ChiS^AF-M^ (liganded) model (see **Fig. S4C** for details). All strains harbored P_chb_-cbp-mCherry and P_const2_-mTFP1 reporter constructs. CBP^Δ^ = the CBP signal sequence translationally fused to mCherry (P_chb_-cbp^1-40^-mCherry). Cells were grown in rich medium (i.e., in the absence of chitin) and fluorescence was determined by plate reader. Data are from three independent biological replicates and shown as the mean ± SD.

**Fig. S8.**
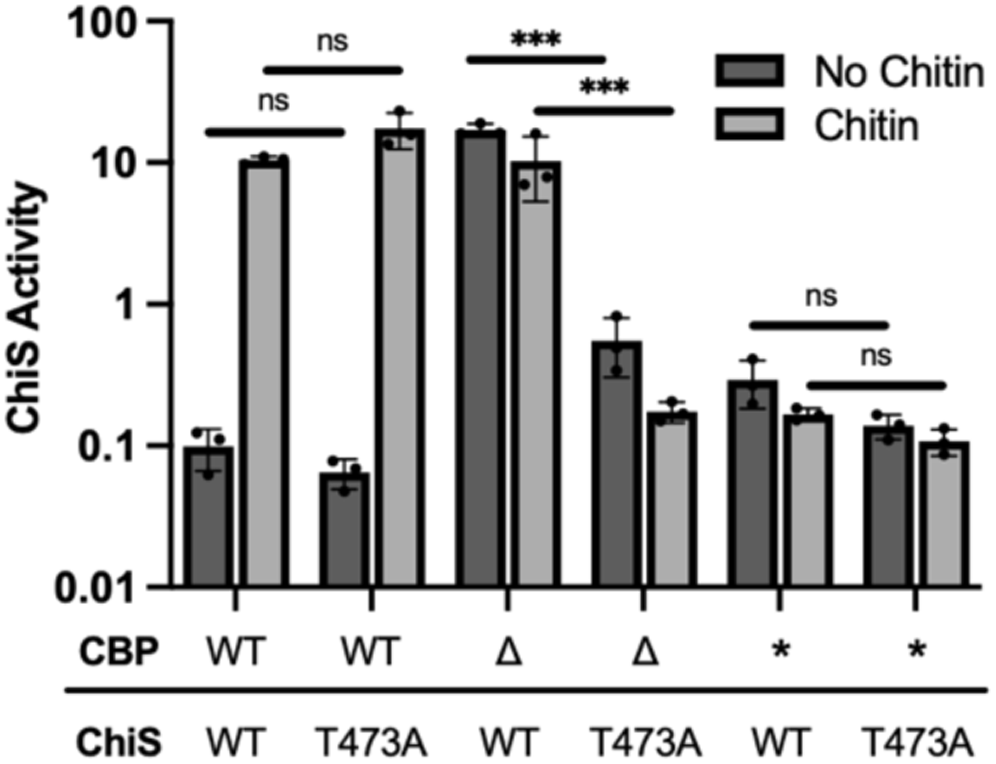
The phosphatase-deficient ChiS^T473A^ allele requires liganded (chitin-bound) CBP for allosteric activation. Transcriptional reporter assays to assess ChiS activity in strains harboring the indicated mutations. All strains harbored P*_chb_-cbp-mCherry* and P*_const2_-mTFP1* reporter constructs. CBP* = CBP^W389A^ ^W539A^, a mutant allele that cannot bind to chitin. CBP^Δ^ = the CBP signal sequence translationally fused to mCherry (P*_chb_-cbp^1-40^-mCherry)*. Cells were grown in either rich medium (No chitin; dark grey bars) or on chitin flakes (chitin; light grey bars) and fluorescence was determined by microscopy. Data are from three independent biological replicates (*n* = 300 cells analyzed per replicate) and shown as mean ± SD. Statistical comparisons were made by one-way ANOVA with Tukey’s multiple comparison test on the log-transformed data. ns = not significant, *** = *p* < 0.001.

**Fig. S9.**
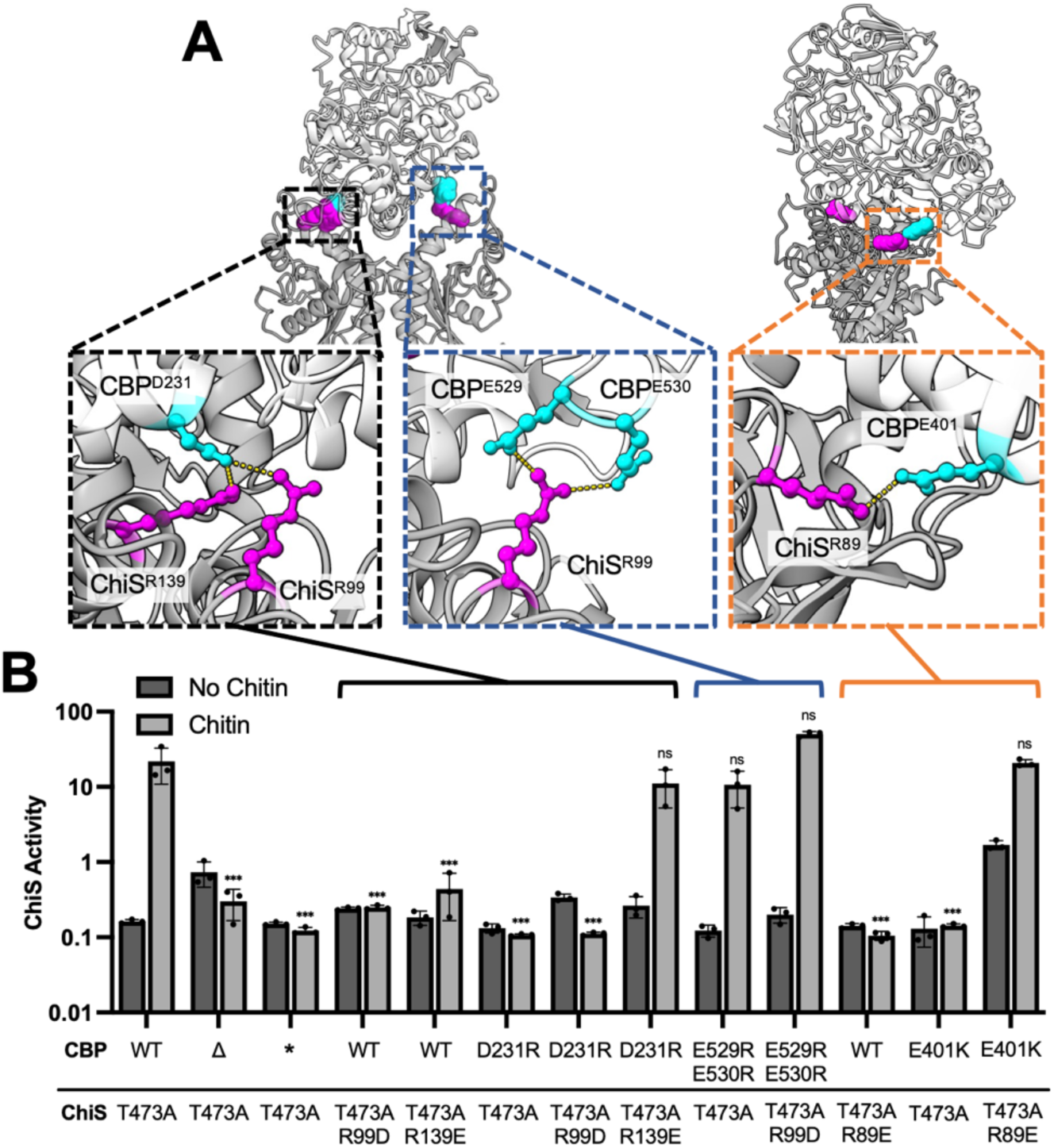
Multiple intermolecular salt bridges specific to CBP-ChiS^AF-M^ (liganded) are necessary for chitin-induced activation. (**A**) The positions of CBP^D231^, CBP^E529,E530^, CBP^E401^, ChiS^R99,R139.^, and ChiS^R89^ are highlighted in the CBP-ChiS^AF-M^ (liganded) model to show the salt bridges formed. CBP is shown in light grey, while ChiS is shown in dark grey. Insets highlight the position of side chains with CBP residues colored cyan, ChiS residues colored magenta, and their associated salt bridges denoted by a yellow dashed line. (**B**) Transcriptional reporter assays to assess ChiS activity in strains harboring the indicated mutations. All strains harbored P*_chb_-cbp-mCherry* and P*_const2_-mTFP1* reporter constructs. CBP* = CBP^W389A,W539A^, a mutant allele that cannot bind to chitin. CBP^Δ^ = the CBP signal sequence translationally fused to mCherry (P*_chb_-cbp^1-40^-mCherry)*. Cells were grown in either rich medium (No chitin; dark grey bars) or on chitin flakes (chitin; light grey bars) and fluorescence was determined by microscopy. The “No Chitin” data for CBP^WT^ ChiS^T473A^, CBP^Δ^ ChiS^T473A^ and CBP^W389A,W539A^ ChiS^WT^ were duplicated from Fig. 4B and included here for ease of comparison. Data are from three independent biological replicates (*n* = 300 cells analyzed per replicate) and shown as the mean ± SD. Statistical comparisons were made by one-way ANOVA with Tukey’s multiple comparison test on the log-transformed data. ns = not significant, *** = *p* < 0.001. Statistical identifiers directly above bars represent comparisons to the parent from the same experimental condition.

**Fig. S10.**
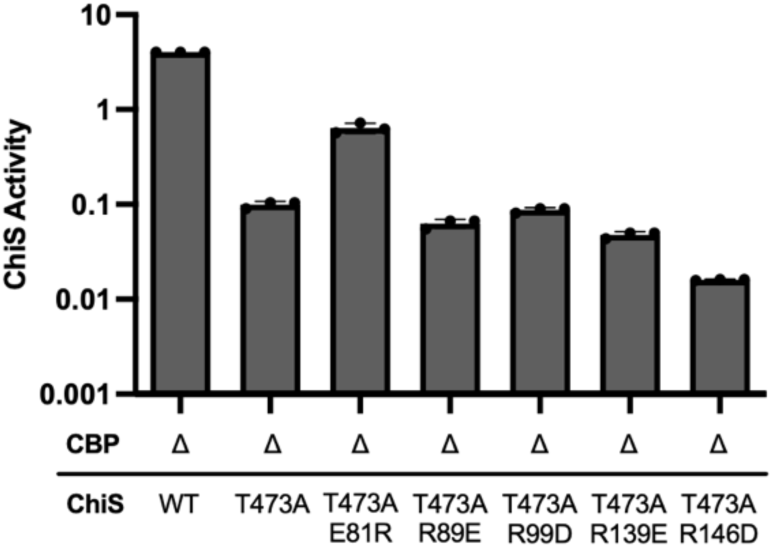
Testing CBP-ChiS^AF-M^ interface mutants for a gain-of-function phenotype. Transcriptional reporter assays to assess ChiS activity in strains harboring the indicated mutations. All strains harbored CBP^Δ^ (P*_chb_-cbp^1-40^-mCherry*) and P*_const2_-mTFP1* reporter constructs. Cells were grown in rich medium (*i.e.*, in the absence of chitin) and fluorescence was determined by plate reader. Data are from three independent biological replicates and shown as mean ± SD. The ChiS^T473A,E81R^ allele exhibited a gain-of-function phenotype, thus, this interaction (**Fig. S4C**) was excluded from further analysis.

**Fig. S11.**
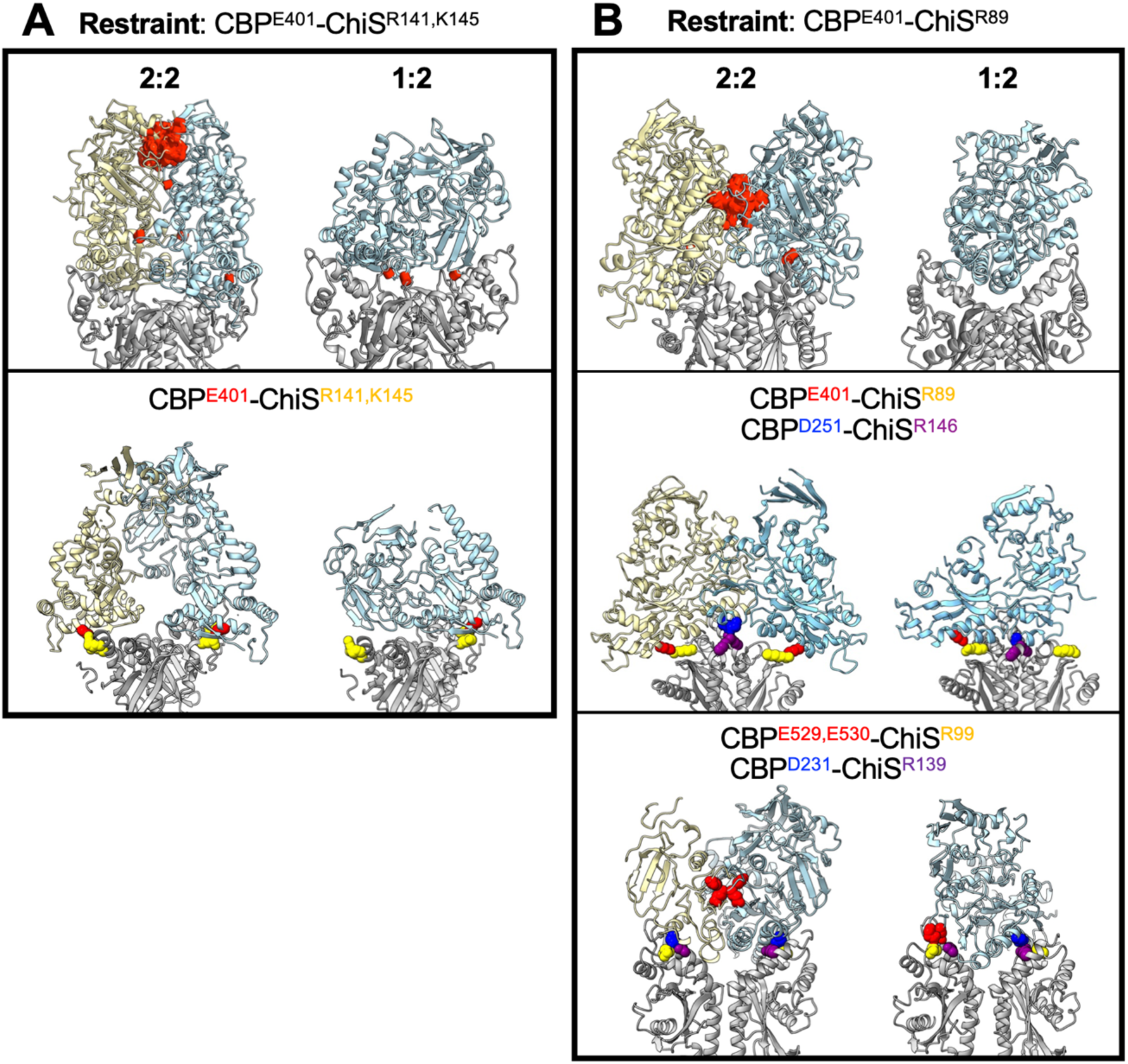
Restrained modelling supports a 1:2 CBP-ChiS stoichiometry. Top ranked models from Chai-1 using 2:2 and 1:2 stoichiometries for CBP (light blue & light yellow) and ChiS (grey) as indicated. Red cylinders on the top models demarcate steric clashes. (**A**) Modeling was restrained using the CBP^E401^-ChiS^R141,K145^ contact that is specific to the unliganded conformation. (**B**) Modeling was restrained using the CBP^E401^-ChiS^R89^ contact that is specific to the liganded conformation. The side chains of the indicated CBP and ChiS residues are shown (color matched to the name in each model) to highlight the spatial proximity of these experimentally validated contacts in each model. Most notably, the CBP^E529,E530^-ChiS^R99^ contact is not maintained in 2:2 models, indicating that the liganded conformation cannot adopt the 2:2 stoichiometry.

**Fig. S12.**
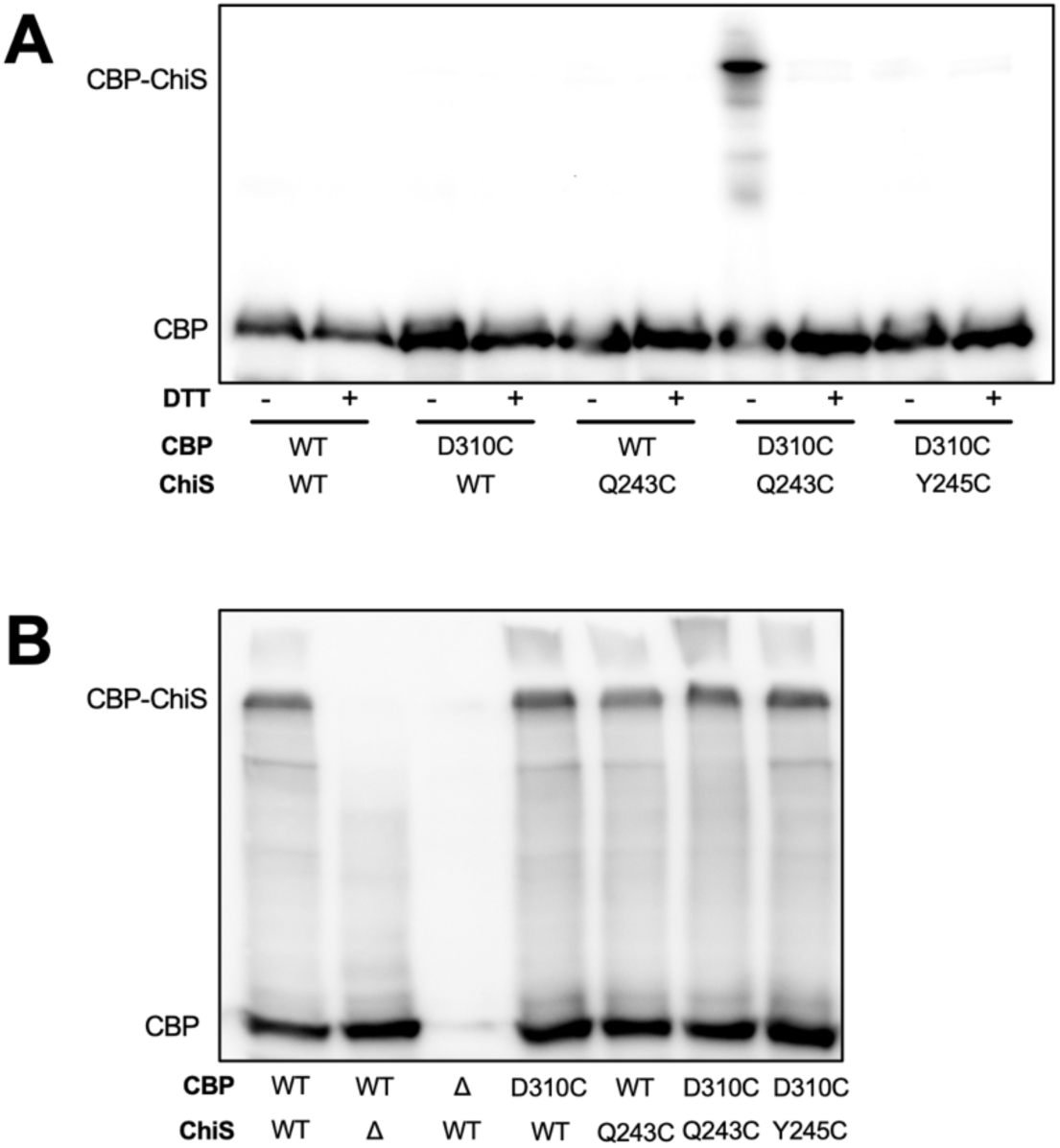
Cysteine knock-ins to CBP-ChiS do not inhibit CBP-ChiS interactions and form a sequence-specific disulfide lock that can be reversed under reducing conditions. Western blots probed using α-mCherry primary antibody. All strains harbored P_tac_-chiS^ΔDBD^-tetR and P_chb_-cbp-mCherry constructs with the indicated mutations and were grown aerobically. (**A**) Non-reducing western blots to assess CBP-ChiS disulfide crosslinking. All strains harbor P_tac_-chiS^ΔDBD^-tetR and P_chb_-cbp-mCherry constructs with the indicated mutations. Lysates were either run directly (-DTT; oxidizing conditions) or treated with DTT to reduced disulfide bonds prior to gel electrophoresis. (**B**) NHS-ester crosslinking with DSS was used to assess CBP-ChiS interactions by Western Blot. Data are representative of two independent biological replicates. The shifted band absent in the *cbp^WT^* Δ*chiS* sample defines the band representing the CBP-ChiS complex. Data are representative of two independent biological replicates.

**Fig. S13.**
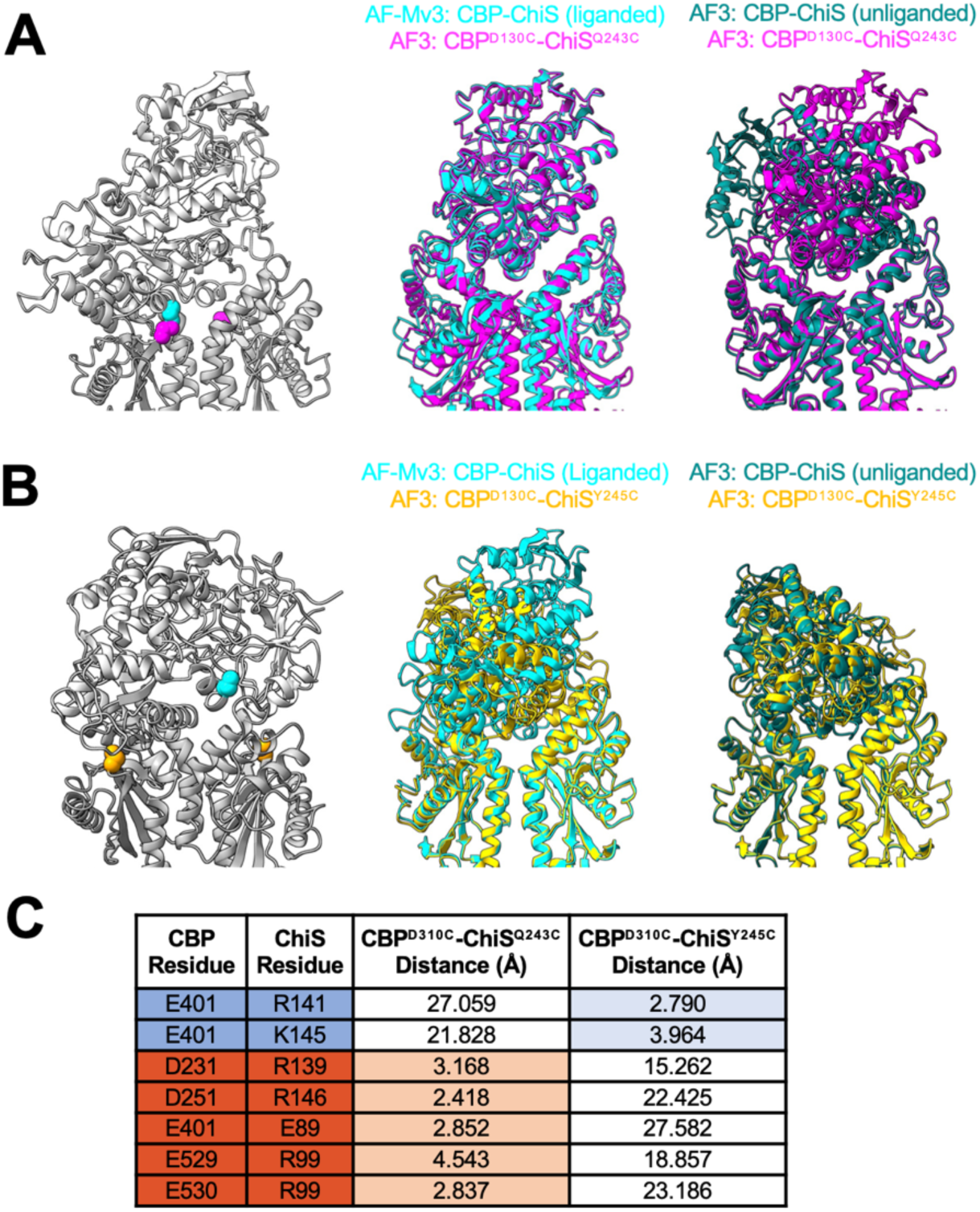
AF3 modeling of the disulfide lock prioritizes the liganded conformation. (**A**) Highlighted CBP^D310C^-ChiS^Q243C^ residues (cyan and magenta, respectively) demonstrates that they are closely juxtaposed in the AF3 model, which is consistent with the liganded CBP-ChiS conformation. CBP is colored light grey, while ChiS is colored dark grey. Overlays of the AF3 CBP^D310C^-ChiS^Q243C^ model (magenta) with CBP-ChiS models from AF-Mv3 (liganded, cyan) and AF3 (unliganded, teal) to highlight the orientation of CBP within each model. (**B**) Highlighted CBP^D310C^-ChiS^Y245C^ residues (cyan and yellow, respectively) are demonstrates that they remain far apart in the AF3 model, which is consistent with the unliganded conformation. CBP is colored light grey, while ChiS is colored dark grey. Overlays of AF3 CBP^D310C^-ChiS^Y245C^ model (yellow) with CBP-ChiS models from AF-Mv3 (liganded, cyan) and AF3 (unliganded, teal) to highlight the orientation of CBP within each model. (**C**) Smallest distance between side chain atoms of the indicated residue pairs are shown.

**Fig. S14.**
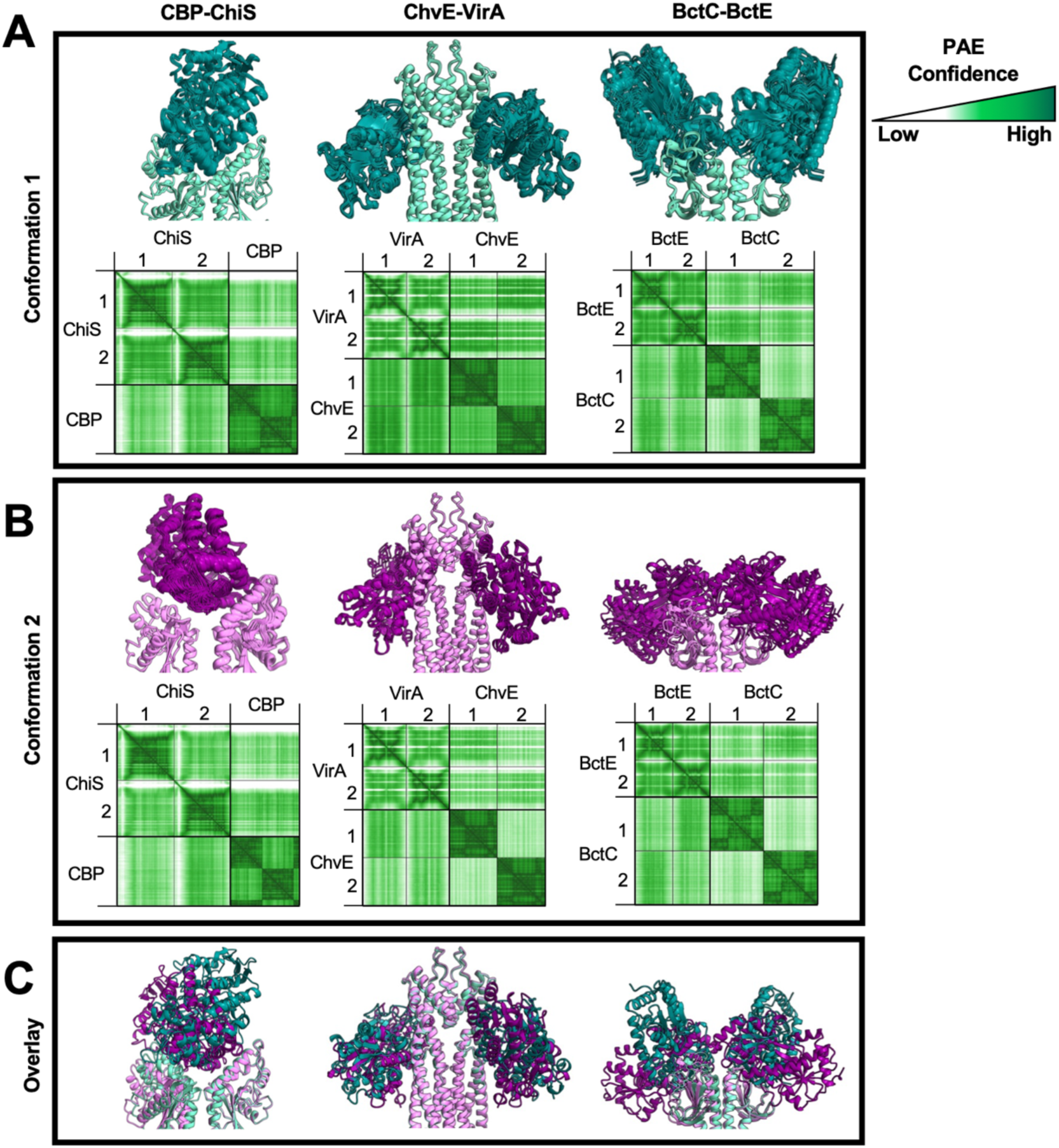
AF-M predicts distinct conformations for other SBP-HK signaling complexes. AF-M reveals two distinct conformations for each SBP-HK pair. Models are aligned using one HK protein chain as the reference point. PAE confidence plots for the top ranked model in each conformation are shown below. (**A**) The top-ranked conformation for CBP-ChiS, ChvE-VirA, and BctC-BctE. Out of 25 independent seeds, this conformation was predicted in 11, 9, and 8 models, respectively. The models that shared this conformation have been aligned to show model-to-model agreement. SBPs are shown in teal and HKs shown in aquamarine. (**B**) The 2^nd^-ranked conformation for CBP-ChiS, ChvE-VirA, and BctC-BctE. Out of 25 independent seeds, this conformation was predicted in 9, 3, and 4 models, respectively. The models that shared this conformation have been aligned to show model-to-model agreement. SBPs shown in purple and HKs shown in violet. (**C**) Alignment of the top-ranked model in each conformation highlights the major differences between the orientation of the SBP in each conformation. Conformation 1 shown in teal (SBP) and aquamarine (HK). Conformation 2 shown in dark magenta (SBP) and violet (HK).

**Fig. S15.**
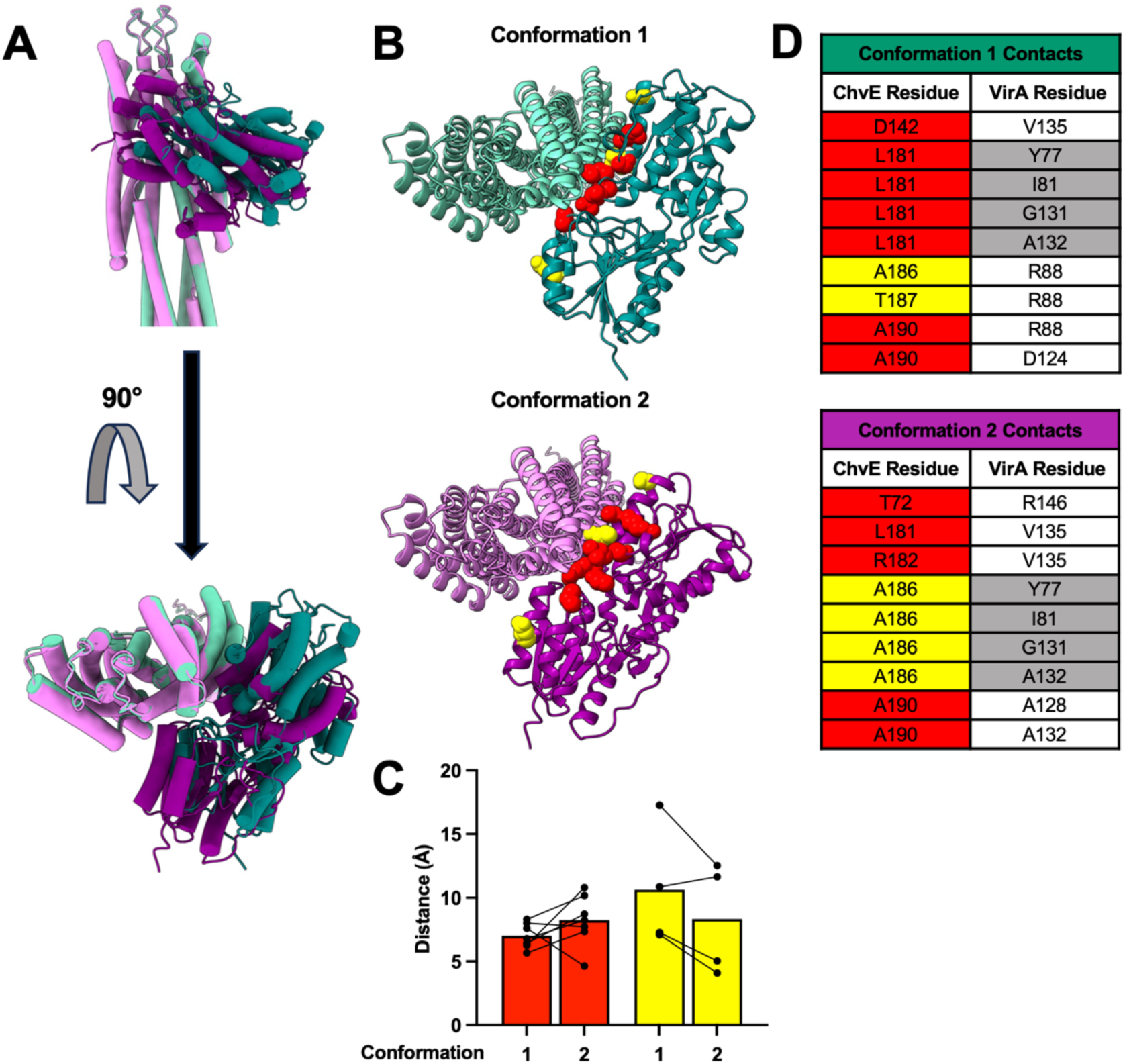
ChvE residues critical for VirA-dependent activity are predicted at the ChvE-VirA interface. (**A**) Cartoon models (alpha-helices shown as cylinders) highlight the large differences in the ChvE-VirA interaction interface between the two predicted conformations of this complex (see **Fig. S9**). For conformation 1, ChvE is colored teal and VirA is colored aquamarine. For conformation 2, ChvE is colored purple and VirA is colored violet. (**B**) Cartoon models highlight the residues previously found to influence ChvE-VirA activity^5^ in both conformations. Residues that, when mutated, decrease VirA activity in the presence of its glucose ligand are colored red. Residues that, when mutated, increase VirA activity in the absence of its glucose ligand are colored yellow. (**C**) The distance between the α-carbons of the indicated ChvE residues (highlighted in **B**) and the closest α-carbon in VirA are plotted for each conformation. (**D**) Predicted contacts for a subset of the ChvE residues shown in **B** highlights residues that undergo partner switching (VirA partner-switch residues are highlighted in grey).

**Movie S1**. A morph between the CBP-ChiS^AF3^ (unliganded) and CBP-ChiS^AF-M^ (liganded) conformations that highlights the large rotational shift of CBP relative to the ChiS dimer. The ChiS dimer is colored yellow and orange, while CBP is colored blue.

**Movie S2**. A morph between the two predicted conformations of ChvE-VirA that highlights the large rotational shift of ChvE relative to VirA. VirA is colored light gray, while ChvE is colored blue. Residues that, when mutated, decrease VirA activity in the presence of its glucose ligand are colored red. Residues that, when mutated, increase VirA activity in the absence of its glucose ligand are colored yellow.

**Movie S3**. A morph between the two predicted conformations of BctC-BctE that highlights the large rotational shift of BctC relative to BctE. BctE is colored light gray, while BctC is colored blue.

**Table S1.**
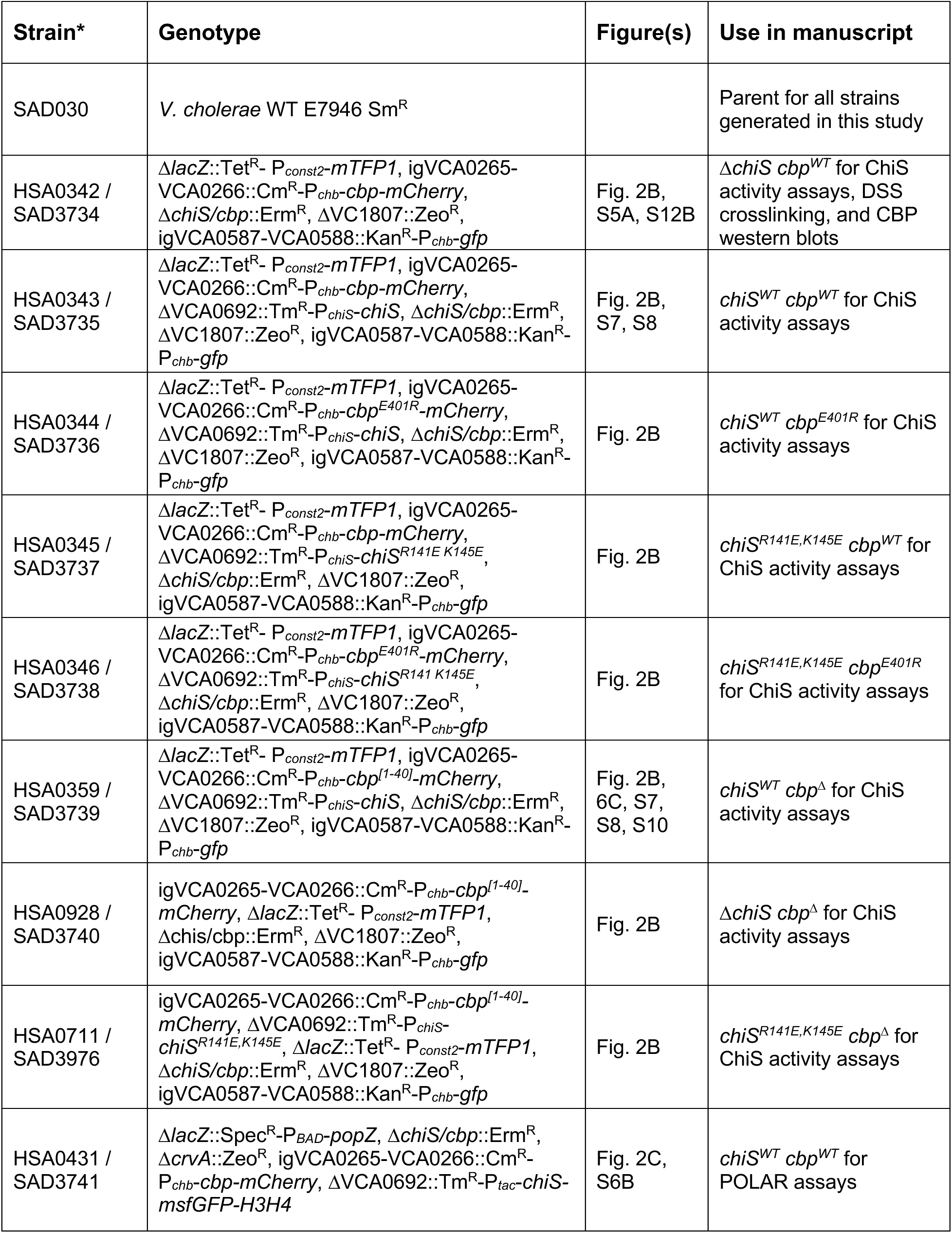
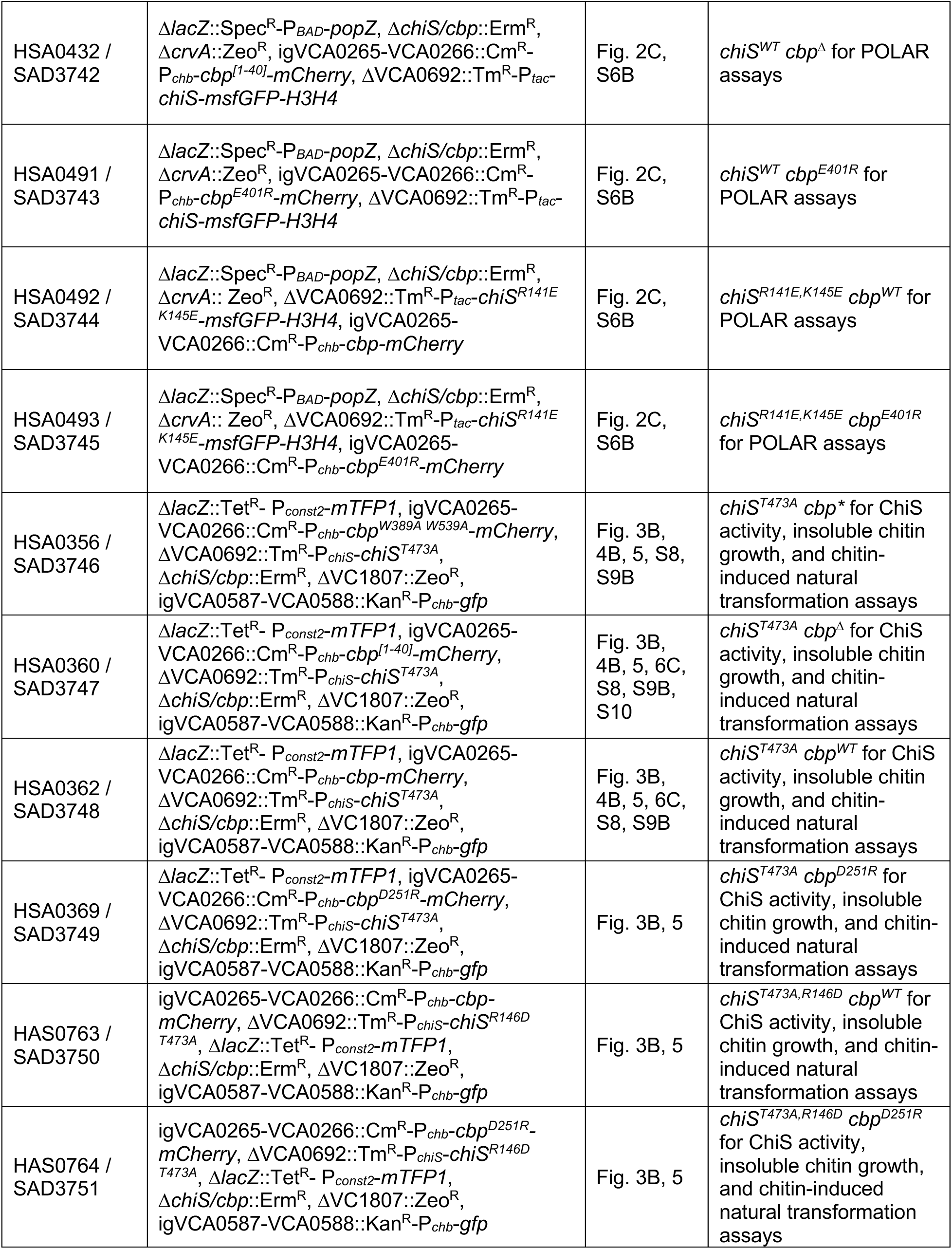
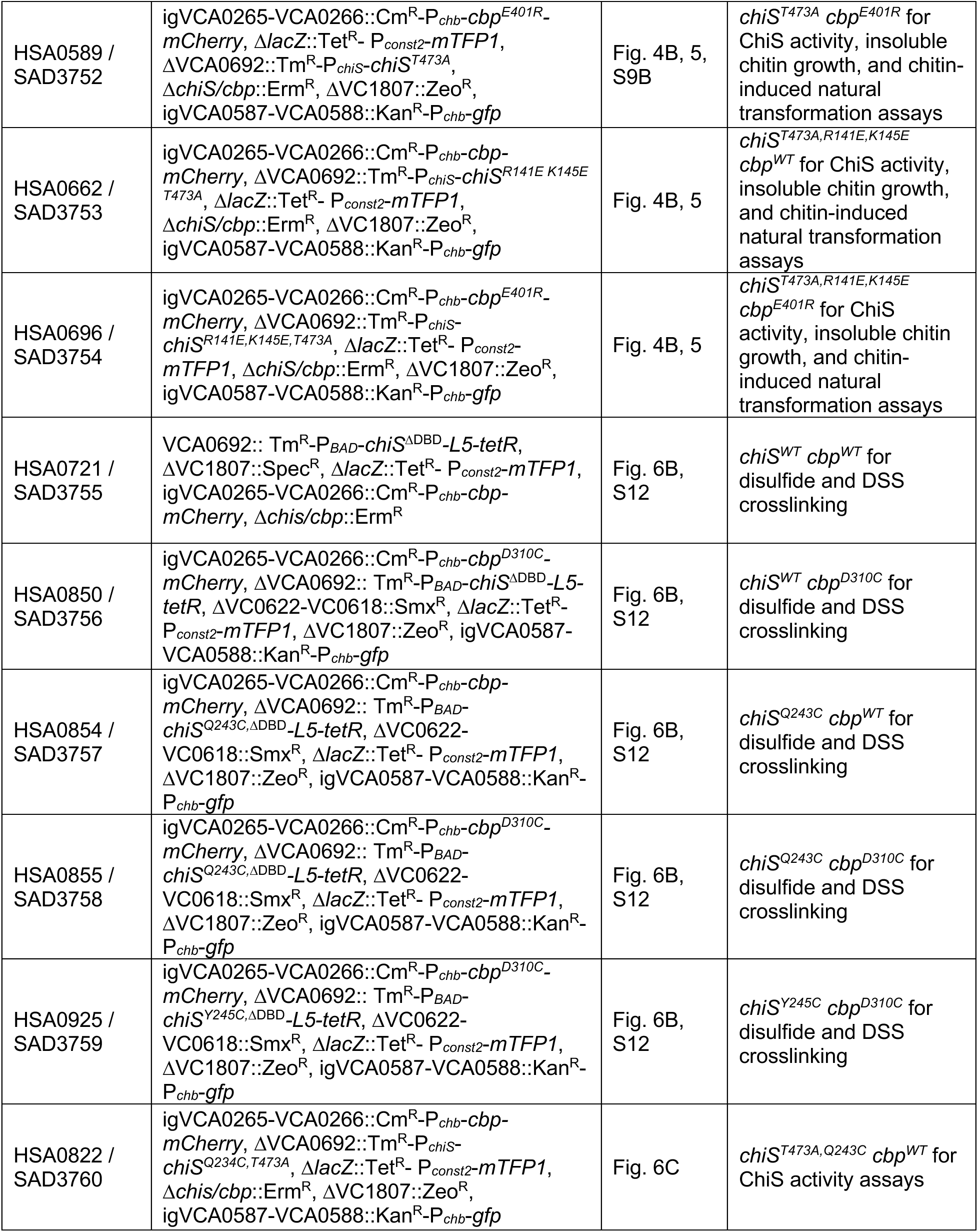
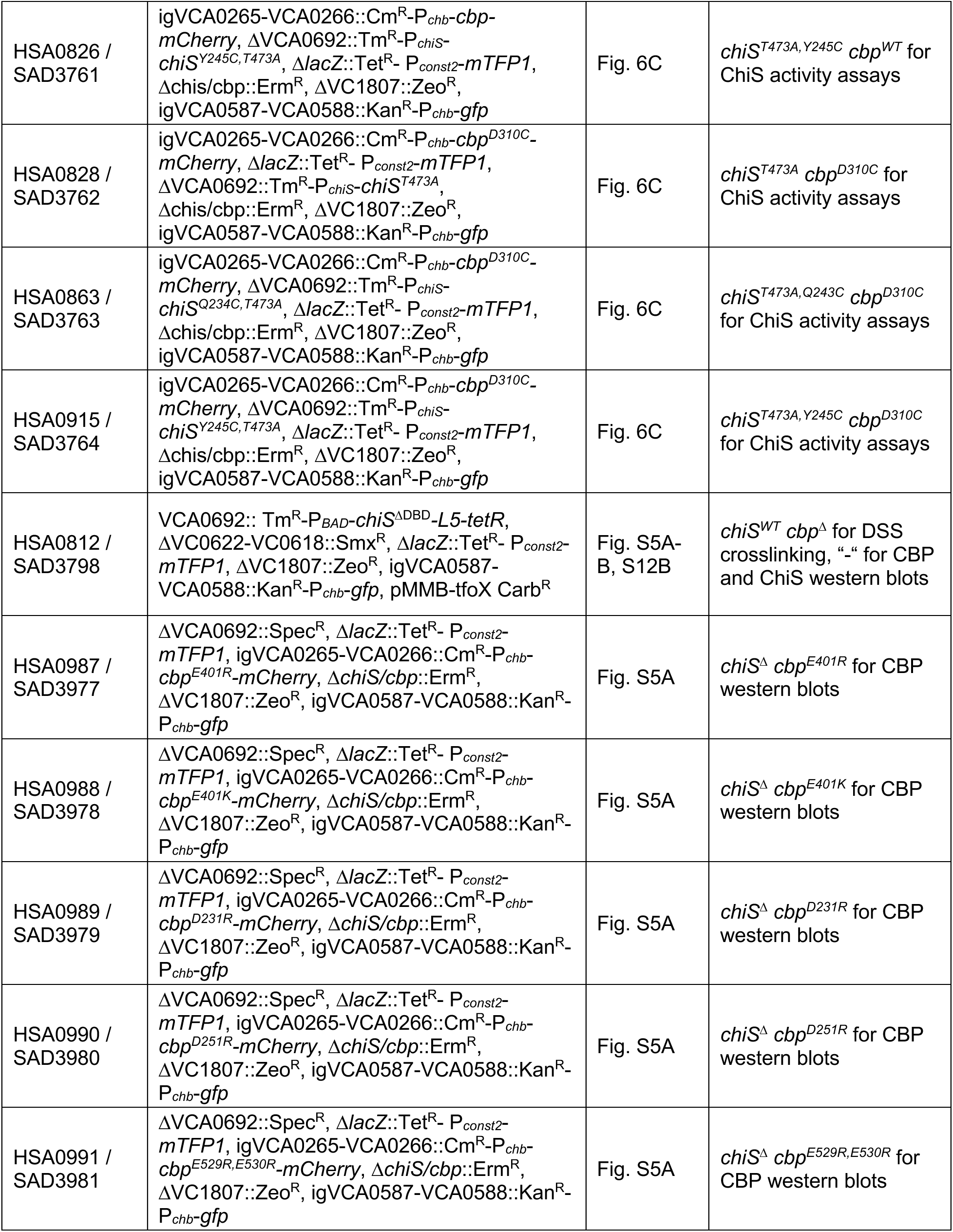
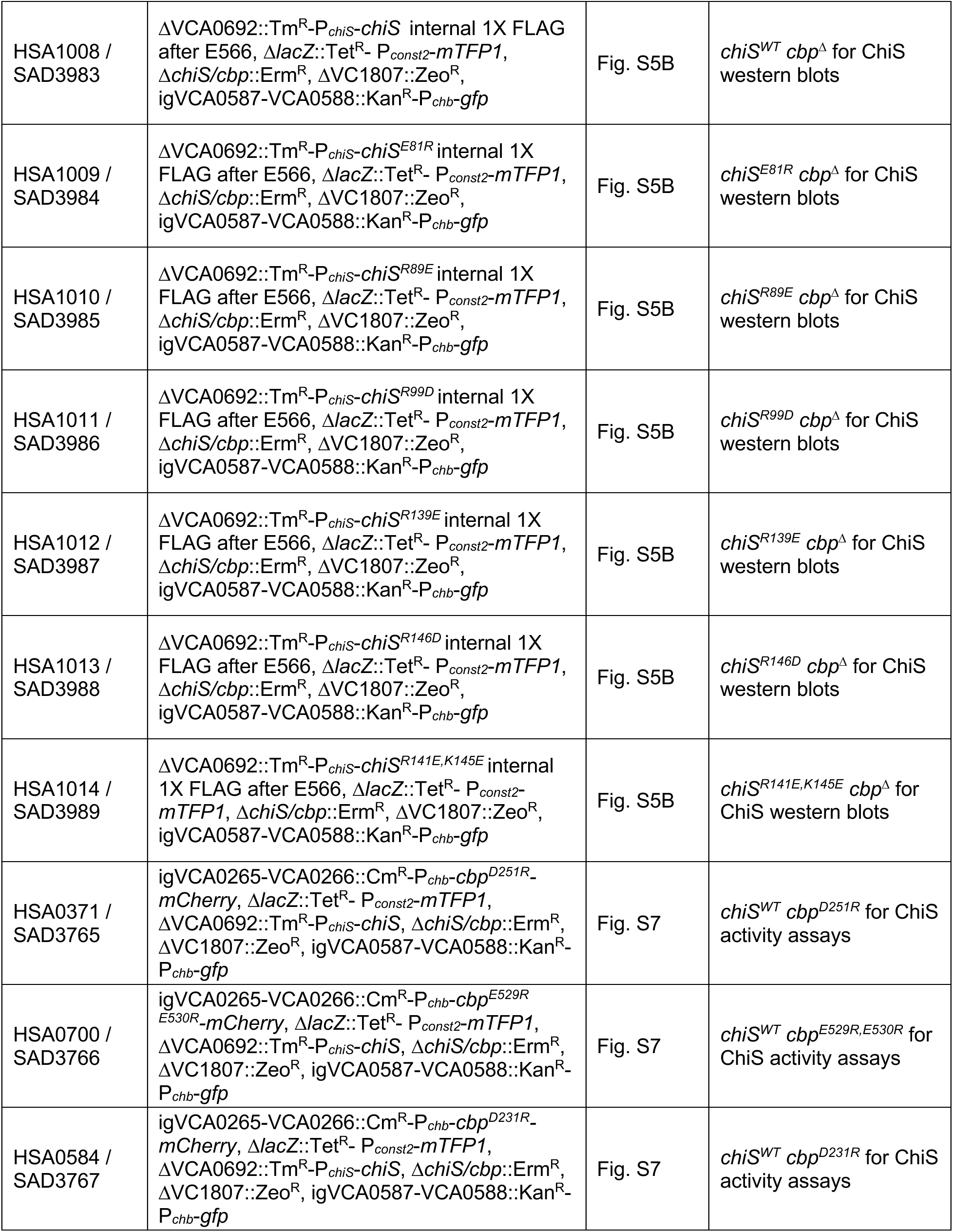
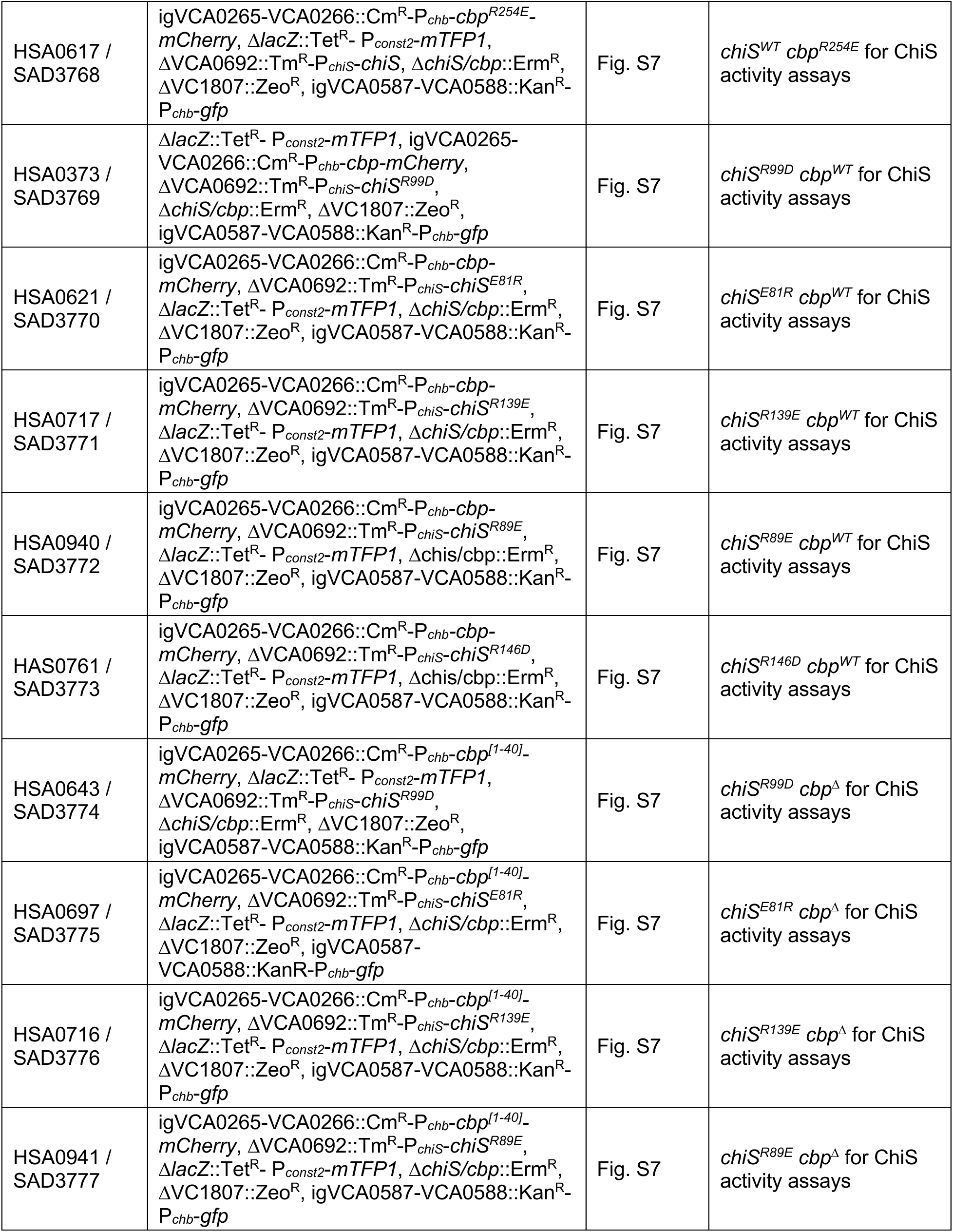
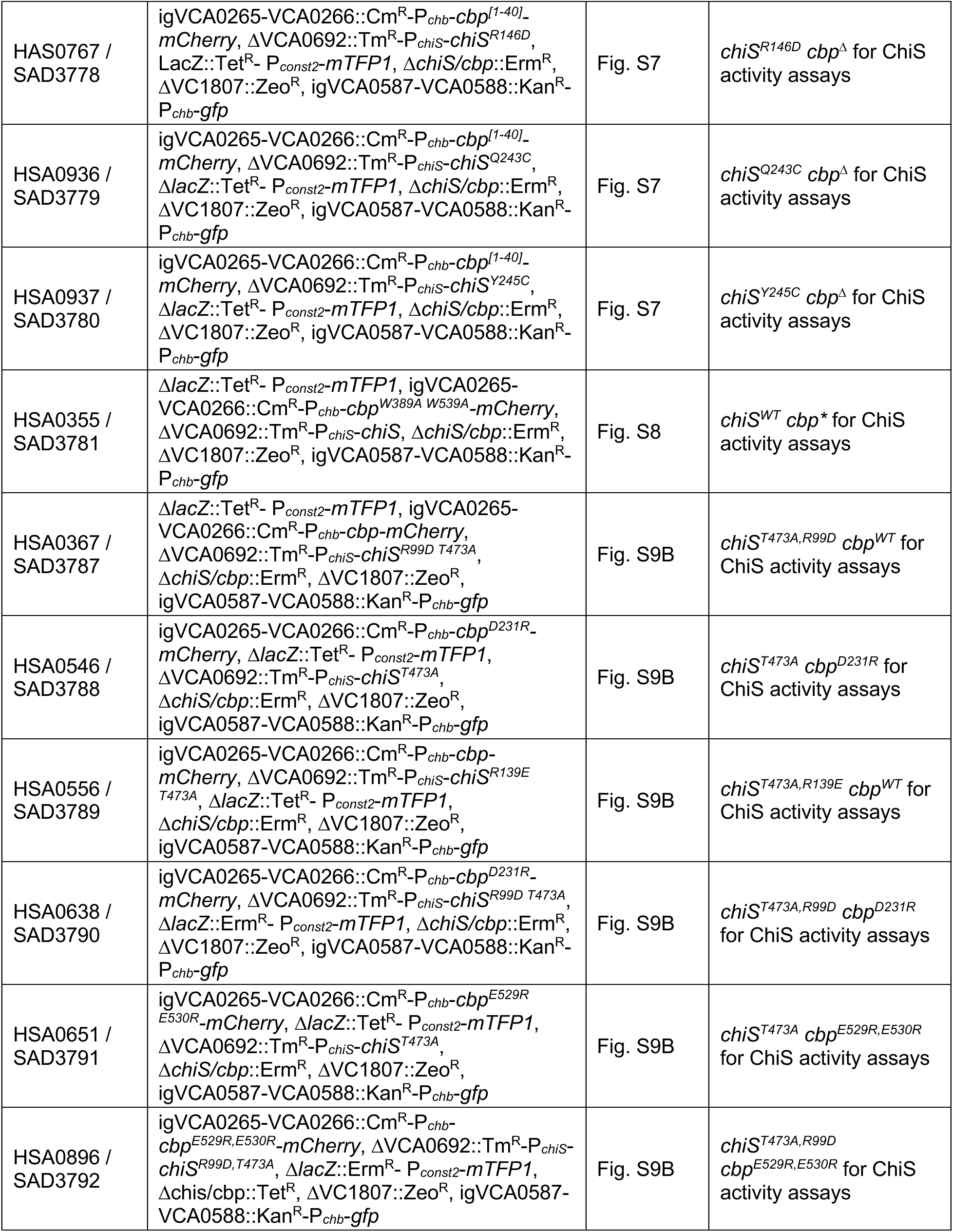
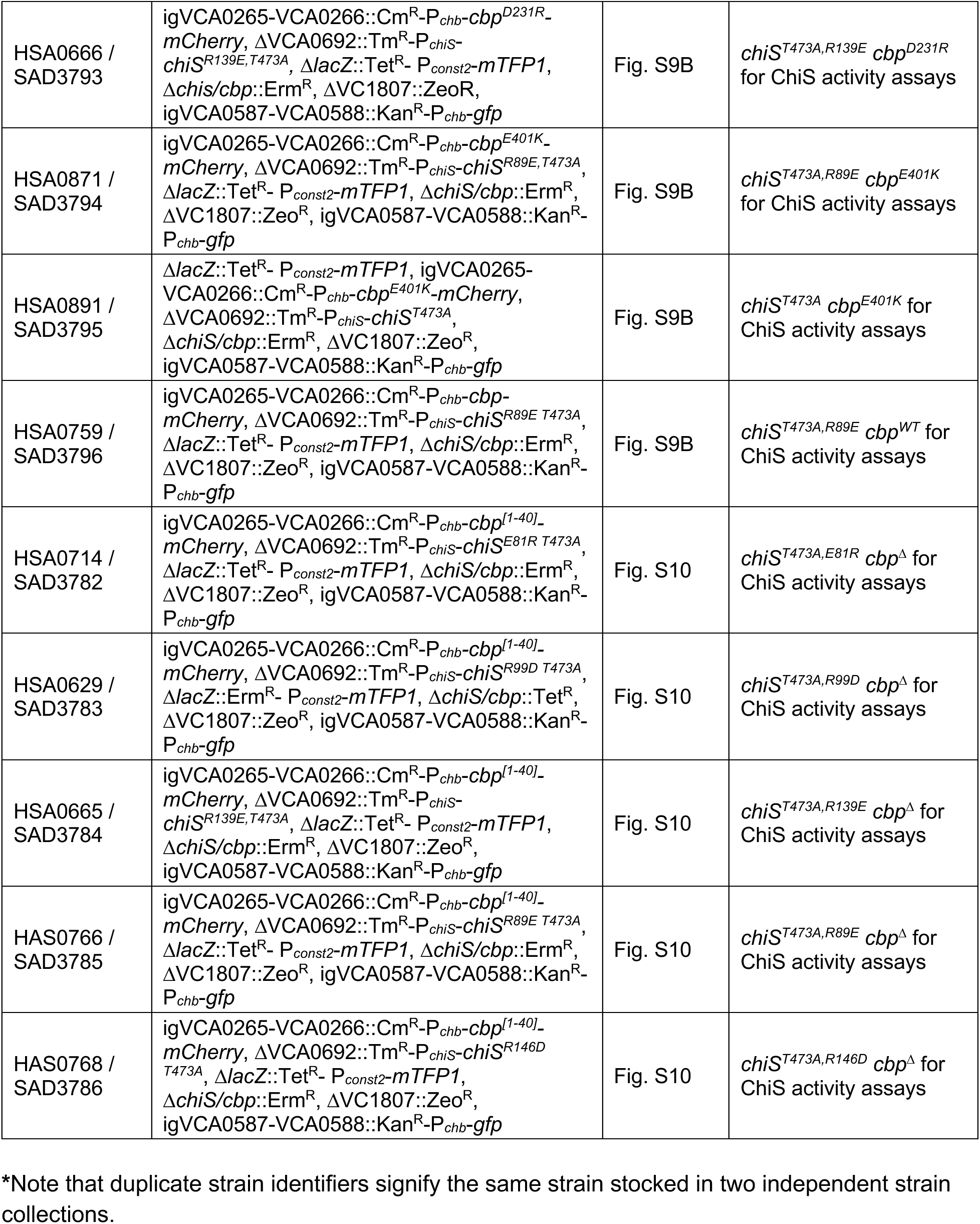
Strains used in this study.

**Table S2.**
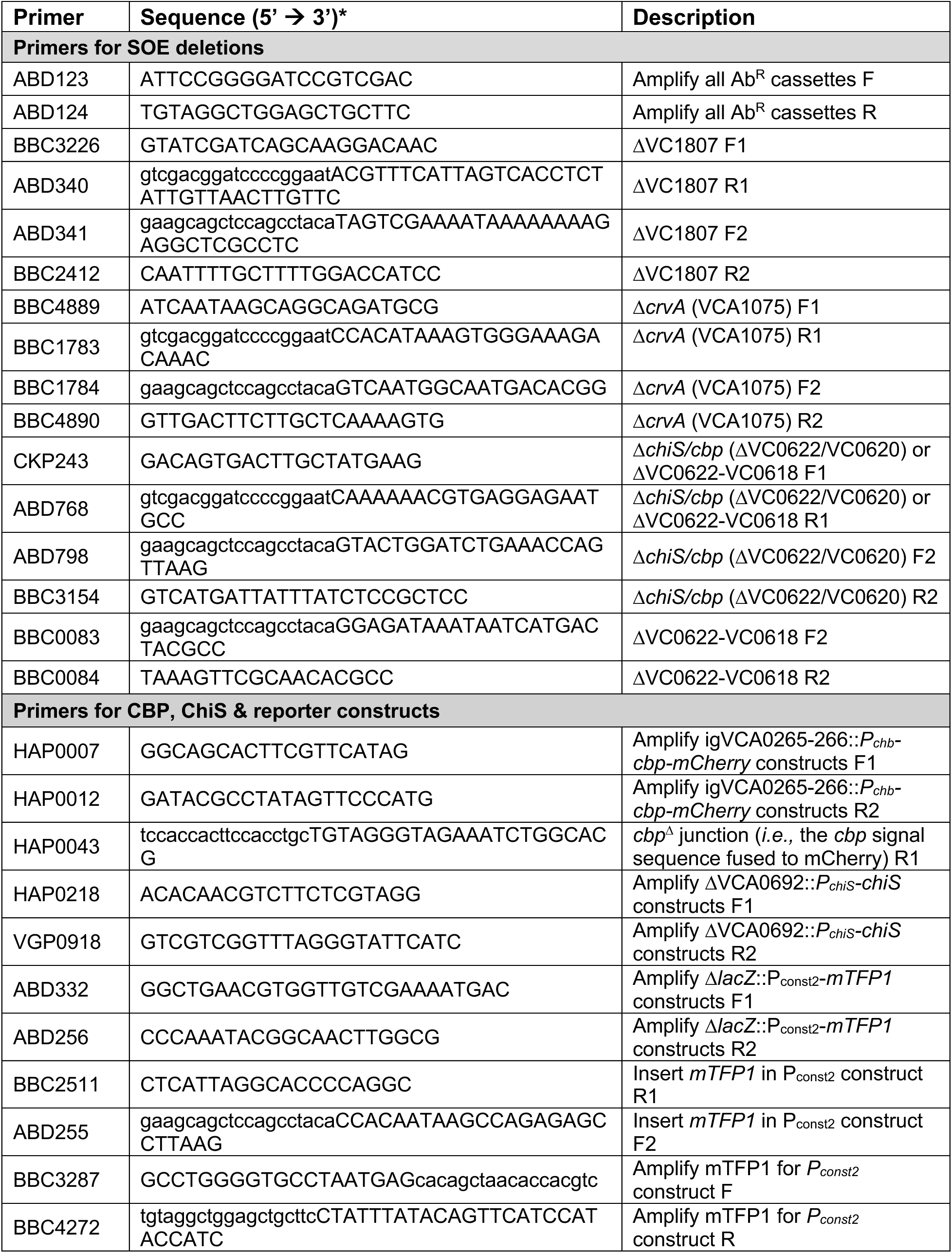
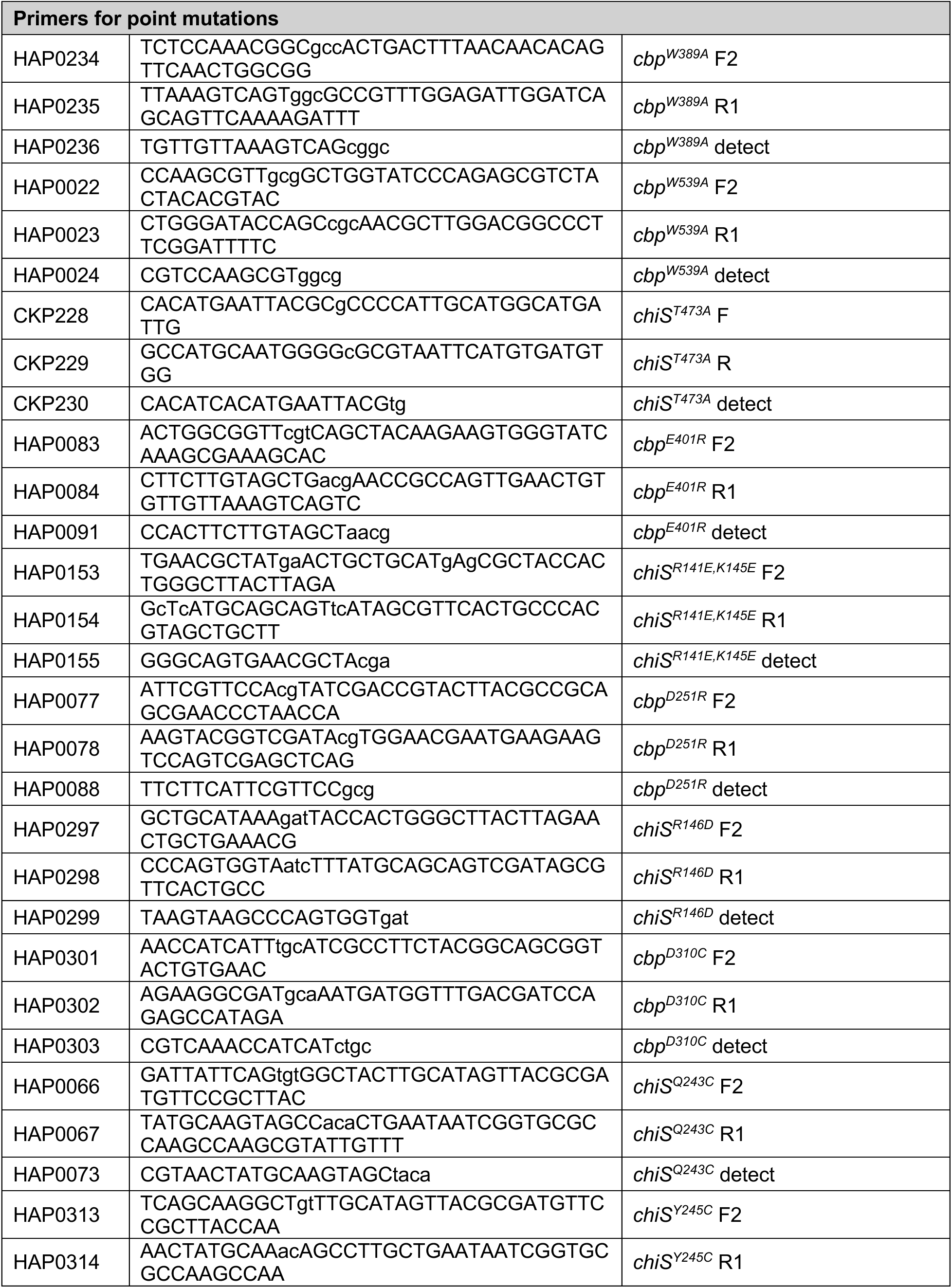
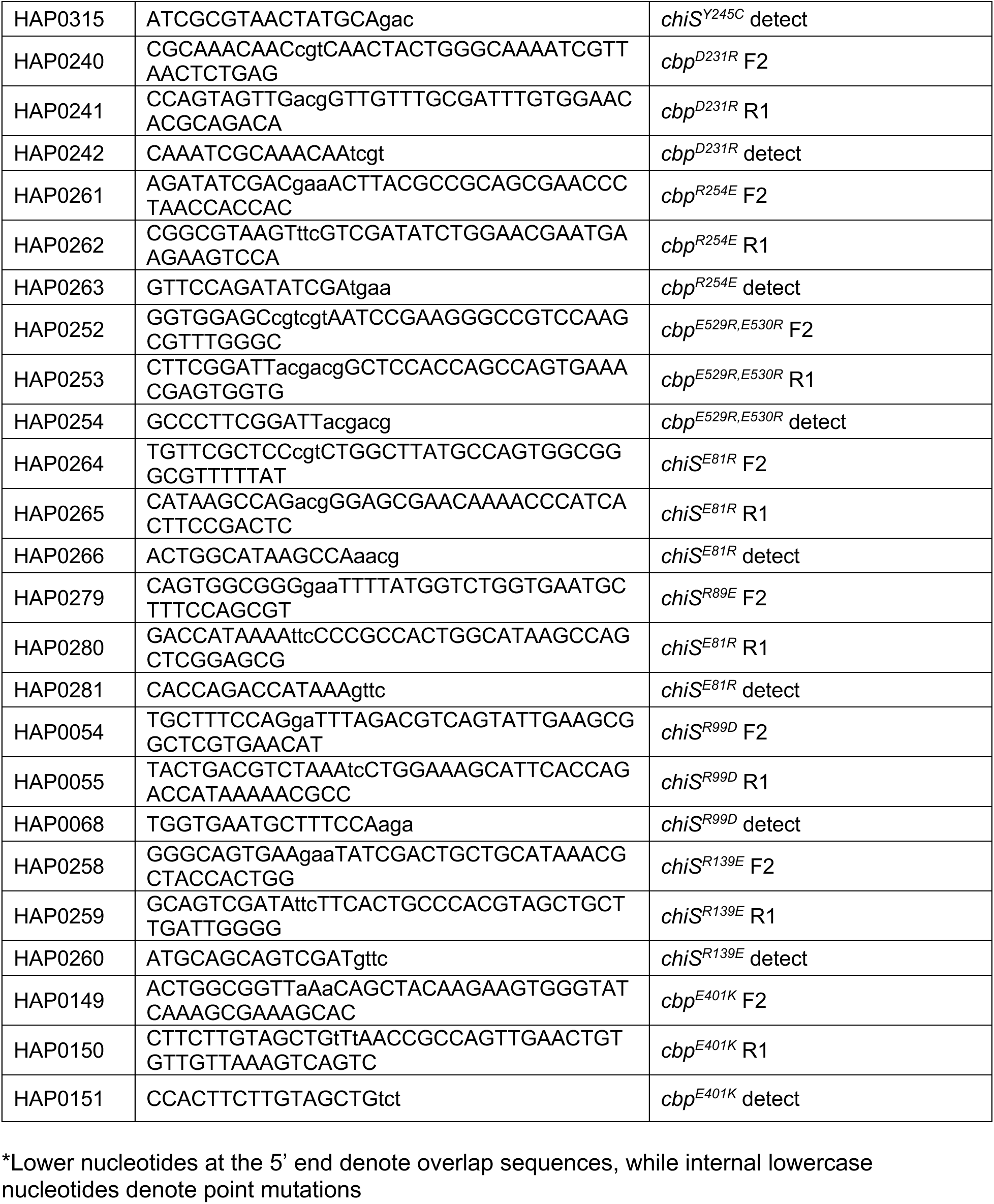
Primers used in this study.

**Dataset S1**. *ChimeraX analysis of contact residues in CBP-ChiS^AF3^ and CBP-ChiS^AF-M^*.

**Dataset S2**. *Summary statistics and statistical comparisons for all quantitative data*.

